# Metabolic disruption impacts tick fitness and microbial relationships

**DOI:** 10.1101/2023.05.26.542501

**Authors:** Sourabh Samaddar, Anya J. O’Neal, Liron Marnin, Agustin Rolandelli, Nisha Singh, Xiaowei Wang, L. Rainer Butler, Parisa Rangghran, Hanna J. Laukaitis, Francy E. Cabrera Paz, Gary M. Fiskum, Brian M. Polster, Joao H. F. Pedra

## Abstract

Arthropod-borne microbes rely on the metabolic state of a host to cycle between evolutionarily distant species. For instance, arthropod tolerance to infection may be due to redistribution of metabolic resources, often leading to microbial transmission to mammals. Conversely, metabolic alterations aids in pathogen elimination in humans, who do not ordinarily harbor arthropod-borne microbes. To ascertain the effect of metabolism on interspecies relationships, we engineered a system to evaluate glycolysis and oxidative phosphorylation in the tick *Ixodes scapularis*. Using a metabolic flux assay, we determined that the rickettsial bacterium *Anaplasma phagocytophilum* and the Lyme disease spirochete *Borrelia burgdorferi*, which are transstadially transmitted in nature, induced glycolysis in ticks. On the other hand, the endosymbiont *Rickettsia buchneri,* which is transovarially maintained, had a minimal effect on *I. scapularis* bioenergetics. Importantly, the metabolite β-aminoisobutyric acid (BAIBA) was elevated during *A. phagocytophilum* infection of tick cells following an unbiased metabolomics approach. Thus, we manipulated the expression of genes associated with the catabolism and anabolism of BAIBA in *I. scapularis* and detected impaired feeding on mammals, reduced bacterial acquisition, and decreased tick survival. Collectively, we reveal the importance of metabolism for tick-microbe relationships and unveil a valuable metabolite for *I. scapularis* fitness.

## Introduction

Arthropod-borne microbes contribute to the global disease burden and are responsible for hundreds of millions of human infections each year^1^. In the United States, the deer tick *Ixodes scapularis* is the predominant arthropod vector and is responsible for transmitting several known human pathogens, including the Lyme disease spirochete *Borrelia burgdorferi* and the obligate intracellular rickettsial bacterium *Anaplasma phagocytophilum* that causes human granulocytic anaplasmosis^2–4^. Although these microbes are best characterized for their ability to cause disease in humans, our knowledge related to the associations between ticks and microbes remains rudimentary. For instance*, I. scapularis* readily acquires *B. burgdorferi* and *A. phagocytophilum* and tolerates their presence transstadially, or throughout developmental stages, but does not transmit to new progeny^5^. On the other hand, *I. scapularis* possesses endosymbiotic bacteria, primarily *Rickettsia buchneri*, that are vertically transmitted from female to progeny but are considered non-pathogenic to humans^6–8^. Overall, tick-microbe relationships are maintained by balancing immune and metabolic responses^9–12^ with fitness advantages conferred by some intracellular bacteria^13–15^.

Parasitism results in a significant fitness cost to the host, such as weight loss, reduced survival or impaired reproductive capacity^16, 17^. Thus, a host may redistribute finite resources upon infection to balance life history programs, including maintenance, growth and reproduction^17–20^. This redistribution, also known as resource allocation, is a main driver in the organism’s response to unfavorable conditions (*e.g.*, pathogen infection) and the push towards maintenance strategies^21, 22^. An example of a maintenance strategy within the cell is metabolic reprogramming, where immune and cancer cells shift their metabolism to aerobic glycolysis, also known as the Warburg effect^23^, to sustain proliferation and/or immune effector functions^22, 24, 25^. Metabolic reprogramming and resource allocation have been largely studied through the lens of evolutionary ecology and mammalian biology^21, 22, 24–30^. However, the metabolic contribution to vector competence remains largely undefined^11, 12, 31^ despite the unique biological relationships between arthropods and the microbes they carry. Previous work suggested that *A. phagocytophilum* and *B. burgdorferi* influence the metabolism of ticks^10, 32–38^. Unfortunately, how disruptions in bioenergetic processes affect ticks on a cellular or systemic level remains fragmented. Furthermore, in what manner ticks respond metabolically to transstadially-infected microbes that cause human diseases compared to a transovarially maintained endosymbiont remain obscure. Finally, individual metabolites that contribute to tick fitness and bacterial acquisition are mostly undetermined.

In this study, we sought to develop a platform for studying bioenergetics in *I. scapularis*. We established a system for manipulating glycolysis and oxidative phosphorylation (OxPhos) in tick cells and measured how disruption in bioenergetic processes affected arthropod fitness. We undertook an unbiased metabolomics approach to characterize metabolic signatures in tick cells infected with either the human pathogen *A. phagocytophilum*^4^ or the endosymbiont *R. buchneri*^6–8^. We identified a key metabolite in ticks, β-aminoisobutyric acid (BAIBA), that is elevated by *A. phagocytophilum*. Finally, we demonstrated that manipulation of gene expression related to the catabolism and anabolism of the D-BAIBA enantiomer in *I. scapularis* influences both tick fitness and bacterial acquisition.

## Results

### Establishment of a system to measure bioenergetics in ticks

Glycolysis, which takes place in the cytoplasm and leads to the production of lactate, and OxPhos, which occurs in the mitochondria and utilizes the electron transport chain (ETC), provides energy for metabolic processes within cells through the generation of adenosine triphosphate (ATP)^22, 24, 25, 30^. Glycolysis and OxPhos can be measured by analyzing the extracellular acidification rate (ECAR) and oxygen consumption rate (OCR), respectively^39^. In the mammalian literature, these processes are evaluated using the Seahorse metabolic flux assay^40^, where cellular metabolism is manipulated by small molecule inhibitors that block enzymatic function. For example, 2-deoxy D-glucose (2-DG) inhibits hexokinase activity, the rate-limiting enzyme in glycolysis, while rotenone, antimycin A, oligomycin, and 2,4- dinitrophenol (2,4-DNP) hinder mitochondrial OxPhos^22, 24, 25, 30^ (Fig. 1A).

**Figure 1:**
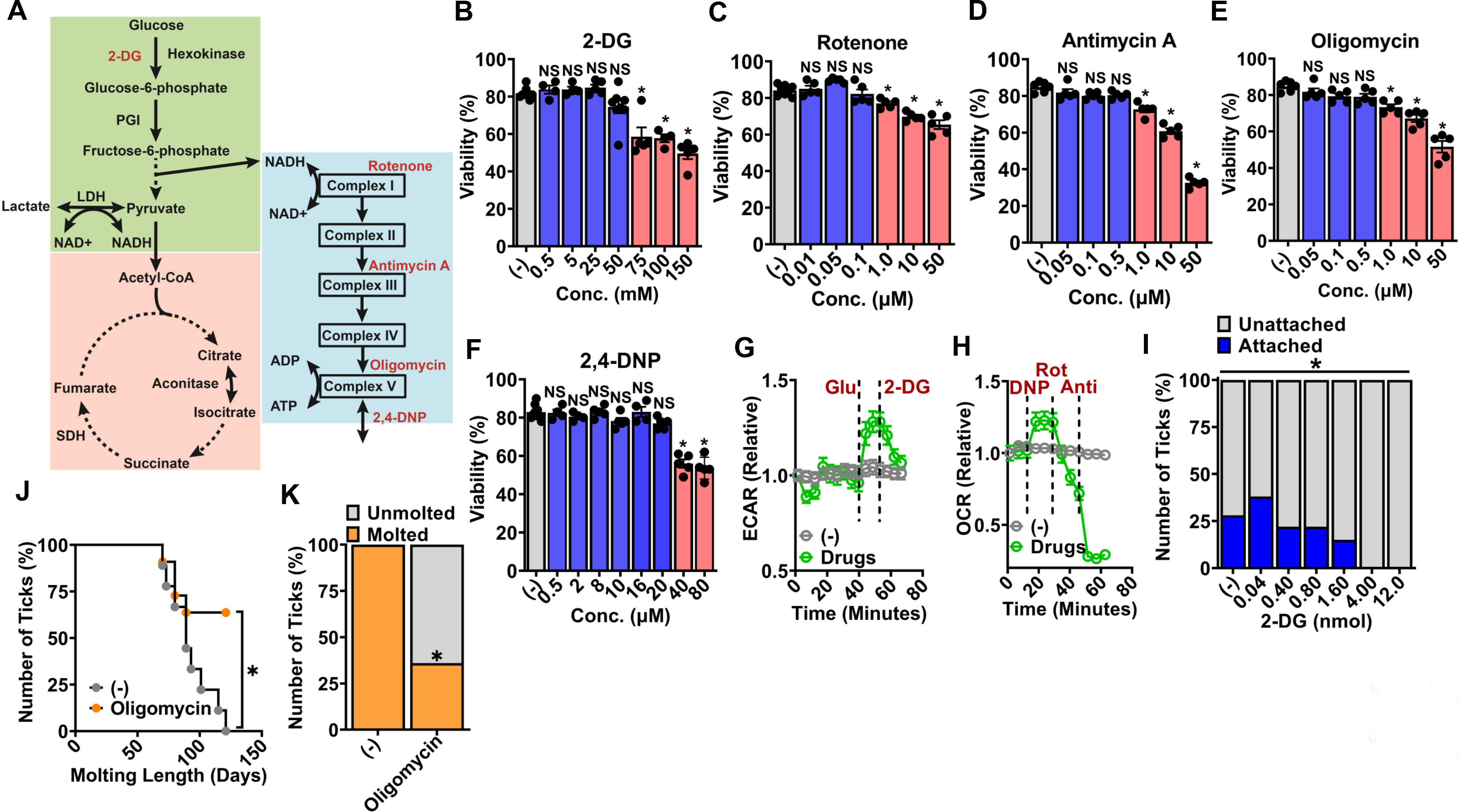
Glycolysis and OxPhos are critical metabolic pathways for fitness programs in ***I. scapularis***. **A.** Abbreviated representation of glycolysis (green), TCA (pink) and OxPhos (blue) with inhibitors that block enzymatic function (red). (**B-F**). Viability measurement of 1x 10^6^ ISE6 cells. Cells were treated with the corresponding inhibitors at indicated concentrations for 48 hours prior to analysis. Data are representative of at least two independent experiments N=4-8. Concentrations that caused a significant decrease in viability are shaded red and non- significant effects are highlighted in blue. **G.** Extracellular acidification rate (ECAR) of 1.2x 10^5^ ISE6 cells treated with chemical inhibitors. Glucose (Glu) and 2-Deoxy-D-Glucose (2-DG) were administered at 25 mM and 50 mM, respectively. Data are representative of at least three independent experiments N=6. **H.** Oxygen consumption rate (OCR) of 1.2x 10^5^ ISE6 cells treated with chemical inhibitors. 2,4-Dinitrophenol (DNP), rotenone and antimycin were administered at 20 µM, 0.1 µM and 0.5 µM, respectively. Data are representative of at least two independent experiments N=6. **I.** Ticks were injected with the respective amounts of 2-DG and placed on C57BL/6 mice overnight (grey). Percentage of ticks that successfully attached on C57BL/6 mice are displayed in blue. Data are representative of at least two independent experiments. Number of ticks used ranged from 25-100 per treatment. **J.** Molting length of ticks following microinjection with oligomycin. Ticks were injected with a sublethal amount of oligomycin (0.8 pmol) prior to feeding. **K.** Percentage of ticks that molted following feeding. Number of ticks used in **J** and **K** ranged from 10-11 per treatment. (**B-F**) One-way ANOVA followed by Dunnett’s test. (**I**) Chi-square test. (**J**) Log rank (Mantel-Cox) test. (**K**) Fisher’s exact test. *, *p*<0.05. Anti = Antimycin A, Rot=rotenone. NS – not significant.

We created a modified version of the commonly used L15C300 medium to culture tick cells (referred to as mL15C)^41^ (table S1), which supports the growth of *I. scapularis* ISE6 cells in the presence of glycolytic and OxPhos molecular inhibitors for at least 48 hours (Figs. 1B-F; fig. S1). Other tick cells available within the scientific community, including IDE12 (*I. scapularis*), AAE2 (*Amblyomma americanum*) and DAE100 (*Dermacentor andersoni*) were not permissive to biochemical manipulation in culture (fig. S2). Thus, we used the *I. scapularis* ISE6 cells for a metabolic flux assay. Using the Seahorse analyzer with drug concentrations that did not affect viability, we demonstrated that ECAR and OCR can be measured in live *I. scapularis* cells (Figs. 1G-H). The addition of glucose led to enhanced extracellular acidification in the mL15C medium containing ISE6 cells (Fig. 1G). The subsequent addition of 2-DG to inhibit hexokinase activity returned extracellular acidification to background levels (Fig. 1G). We also observed an increase in the oxygen consumption rate within *I. scapularis* ISE6 cells upon adding the 2,4- DNP uncoupler. The 2,4-DNP uncoupler disrupts the electrochemical gradient across the mitochondrial inner membrane (Fig. 1H). We decreased the electron transport flow in mitochondria with rotenone (Complex I inhibitor) and antimycin A (Complex III inhibitor), which resulted in reduction of cellular respiration (Fig. 1H). Altogether, we were able to measure critical bioenergetic functions in tick cells, including glycolysis and mitochondrial respiration.

In a tick life cycle, growth is influenced by the uptake of a blood meal and can be directly measured by weight, which is dependent on attachment to mammals. Alternatively, maintenance can be assessed by tick survival and molting to subsequent developmental stages. To determine how metabolic inhibitors influenced ticks *in vivo*, we then injected nymphs with distinct amounts of 2-DG or oligomycin and measured fitness parameters, including attachment, feeding, survival, and molting. For the 2-DG treatment, we observed a dose-dependent effect on tick attachment, but we did not detect differences in other aspects of tick fitness (Fig. 1I and fig. S3). Conversely, elevated concentrations of oligomycin reduced survival in unfed nymphs (fig. S4A). Based on this information, we used an intermediate oligomycin amount (0.8 pmol) in subsequent experiments to measure fitness parameters in fed ticks.

Interestingly, we observed a significant reduction in the molting capacity of ticks injected with oligomycin. All *I. scapularis* injected with the vehicle control phosphate-buffered saline (PBS) molted to adults within 120 days post-feeding on mice (Figs. 1J-K). On the other hand, only 36% of nymphs injected with oligomycin molted during this time period (Figs. 1J-K). Chemical inhibition of the OxPhos complex V by oligomycin did not impair attachment, weight, or survival in fed nymphs (figs. S4B-D). Collectively, our results indicated that oligomycin has distinct effects on fed versus unfed *I. scapularis* nymphs. Furthermore, inhibitors of glycolysis and OxPhos altered biological programs associated with tick growth and maintenance.

### *A. phagocytophilum* and *B. burgdorferi* increase glycolysis in tick cells compared to the endosymbiont *R. buchneri*

During an infection, the host switches its metabolism to glycolysis to fuel immune cells and respond to cellular stress^42, 43^. Conversely, pathogenic microbes upregulate glycolysis to establish infection and turn on virulence programs^22, 42^. We aimed to characterize the tick metabolic response during microbial stimulation. *I. scapularis* may carry *A. phagocytophilum* and *B. burgdorferi*, two bacterial pathogens that are transstadially transmitted^2–4^. Furthermore, they harbor the endosymbiont *R. buchneri*, which is non-pathogenic to humans and passed transovarially to other ticks^6^. Thus, we measured glycolysis (ECAR) and OxPhos (OCR) in tick cells infected with *A. phagocytophilum*, *B. burgdorferi*, or *R. buchneri* (Fig. 2A). After 48 hours of culturing *I. scapularis* ISE6 cells in the mL15C medium, *A. phagocytophilum* or *B. burgdorferi*, but not *R. buchneri*, upregulated glycolysis in a manner consistent with the multiplicity of infection (MOI) (Fig. 2B-D). This glycolytic effect on tick cells was more pronounced when glucose was added to the medium culture (Fig. 2B-D). Importantly, we did not observe any noteworthy impact of bacterial MOI for OxPhos across all conditions based on the OCR analysis (Fig. 2E-G).

**Figure 2:**
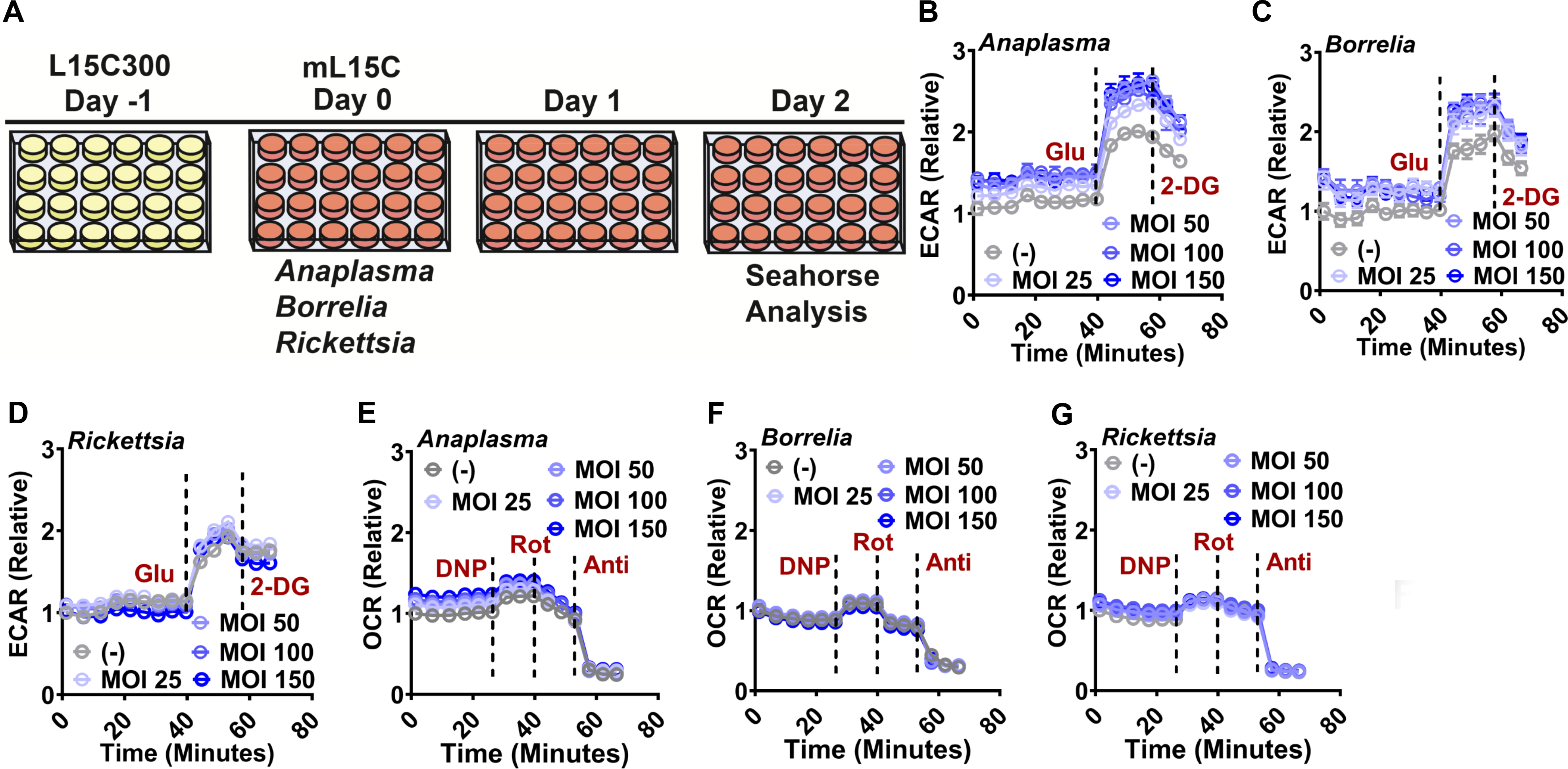
***A. phagocytophilum* and *B. burgdorferi* induce glycolysis upon infection of tick cells. A.** Schematics of infection assay. *I. scapularis* ISE6 cells were cultured in L15C300 medium. At day 0, the L15C300 medium was replaced with the mL15C medium for cell culture. Tick cells were infected with distinct microbial agents at indicated MOI and the Seahorse analysis was done at day 2 post-infection. **B-D.** Extracellular acidification rate (ECAR) of 1.2 x10^5^ ISE6 cells stimulated with (**B**) *A. phagocytophilum*, (**C**) *B. burgdorferi*, or (**D**) *R. buchneri*. Data normalized to unstimulated cells. Data are representative of three independent experiments N=6. **E-G.** Oxygen consumption rate (OCR) of 1.2 x10^5^ ISE6 cells stimulated with (**E**) *A. phagocytophilum*, (**F**) *B. burgdorferi*, or (**G**) *R. buchneri*. Data normalized to unstimulated cells. Data are representative of at least three independent experiments N=6. MOI=multiplicity of infection. Glu=Glucose, 2-DG=2-deoxy-glucose, DNP=2,4-dinitrophenol, Rot=rotenone, Anti=antimycin. DNP, Rot and Anti were administered at 20 µM, 0.1 µM and 0.5 µM, respectively. Glu and 2-DG were administered at 25 mM and 50 mM, respectively.

We then validated the Seahorse metabolic flux assay through colorimetric assays (Fig. 3A). Infection of tick cells with the human pathogens *A. phagocytophilum* or *B. burgdorferi* resulted in increased activity of the enzymes phosphoglucose isomerase (PGI) and lactate dehydrogenase (LDH) (Fig. 3B and 3C, 3E and 3F). Similarly, we observed higher concentrations of lactate after 1 and 24 hours and decreased concentrations of NADH at 24 hours (Fig. 3H and 3I, 3K and 3L). On the other hand, infection of tick cells with *R. buchneri* did not result in significant changes in glycolytic enzymatic activity (Fig. 3D and 3G), although we measured slightly increased levels of lactate and decreased NADH at 24 hours, respectively (Fig. 3J and 3M).

**Figure 3:**
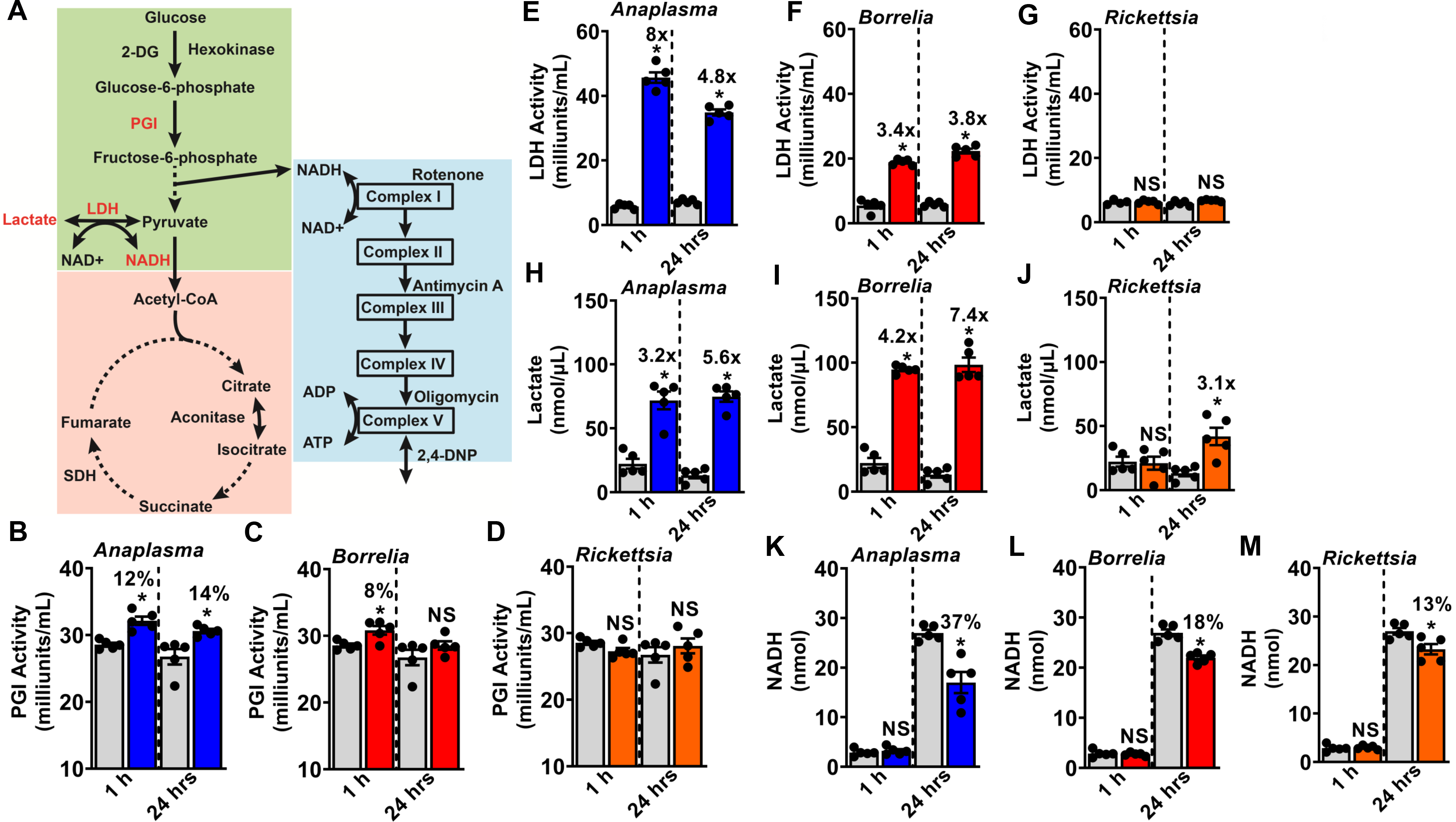
***A. phagocytophilum* and *B. burgdorferi* enhance the glycolytic flux from glucose to lactate in tick cells. A.** Schematic representation of glycolysis (green), TCA (pink) and OxPhos (blue). Readouts for the glycolytic flux to lactate in tick ISE6 cells are highlighted in red. (**B-M**) 1 x10^6^ ISE6 cells were stimulated with *A. phagocytophilum* at a multiplicity of infection (MOI 50) (blue), *B. burgdorferi* (MOI 50) (red), *R. buchneri* (MOI 50) (orange) or left unstimulated (grey) for 1 or 24 hours. (**B-D**) Phosphoglucoisomerase (PGI) and (**E-G**) lactate dehydrogenase (LDH) activity, (**H-J**) lactate and (**K-M**) nicotinamide adenine dinucleotide (NADH) measurements through colorimetric assays. Data represent at least two independent experiments N=5. Statistical significance was evaluated by the unpaired t test with Welch’s correction. *, *p*<0.05. NS – not significant.

We did not observe metabolic changes in components of the tricarboxylic acid (TCA) cycle upon *A. phagocytophilum* infection, as measured by the enzymatic activities of aconitase and succinate dehydrogenase and the metabolites citrate and succinate, respectively (fig. S5). These findings suggested that interactions between tick cells and the human pathogens *A. phagocytophilum* and *B. burgdorferi* affect glycolysis in *I. scapularis* cells. Furthermore, bacterial associations occurring upon *A. phagocytophilum* and *B. burgdorferi* infection of tick cells are distinct from those of the endosymbiont *R. buchneri*.

### Infection with the human pathogen *A. phagocytophilum* alters the metabolism of tick cells compared to the endosymbiont *R. buchneri*

We then tested whether distinct metabolic states (*e.g.*, glycolysis or OxPhos) affected bacterial burden in tick cells. We pre-treated *I. scapularis* ISE6 cells cultured in the mL15C medium with oligomycin, 2,4-DNP, rotenone, antimycin A or 2-DG and measured microbial infection 48 hours later (Fig. 4A). We observed that inhibiting glycolysis with 2-DG had no significant effect on bacterial burden in ISE6 cells (Figs. 4B and 4C). However, impairing OxPhos with oligomycin, rotenone or antimycin A led to a pronounced increase in *A. phagocytophilum* infection of tick cells (Fig. 4B). Comparatively, only a modest increase in *R. buchneri* infection was noted after blocking OxPhos (Fig. 4C). Importantly, treatment of tick cells with the uncoupler 2,4-DNP neither affected *A. phagocytophilum* nor *R. buchneri* infection (Fig. 4B and 4C). Overall, these results indicated that impeding OxPhos in tick cells created an environment that benefited *A. phagocytophilum*, and to a lesser extent, *R. buchneri*.

**Figure 4:**
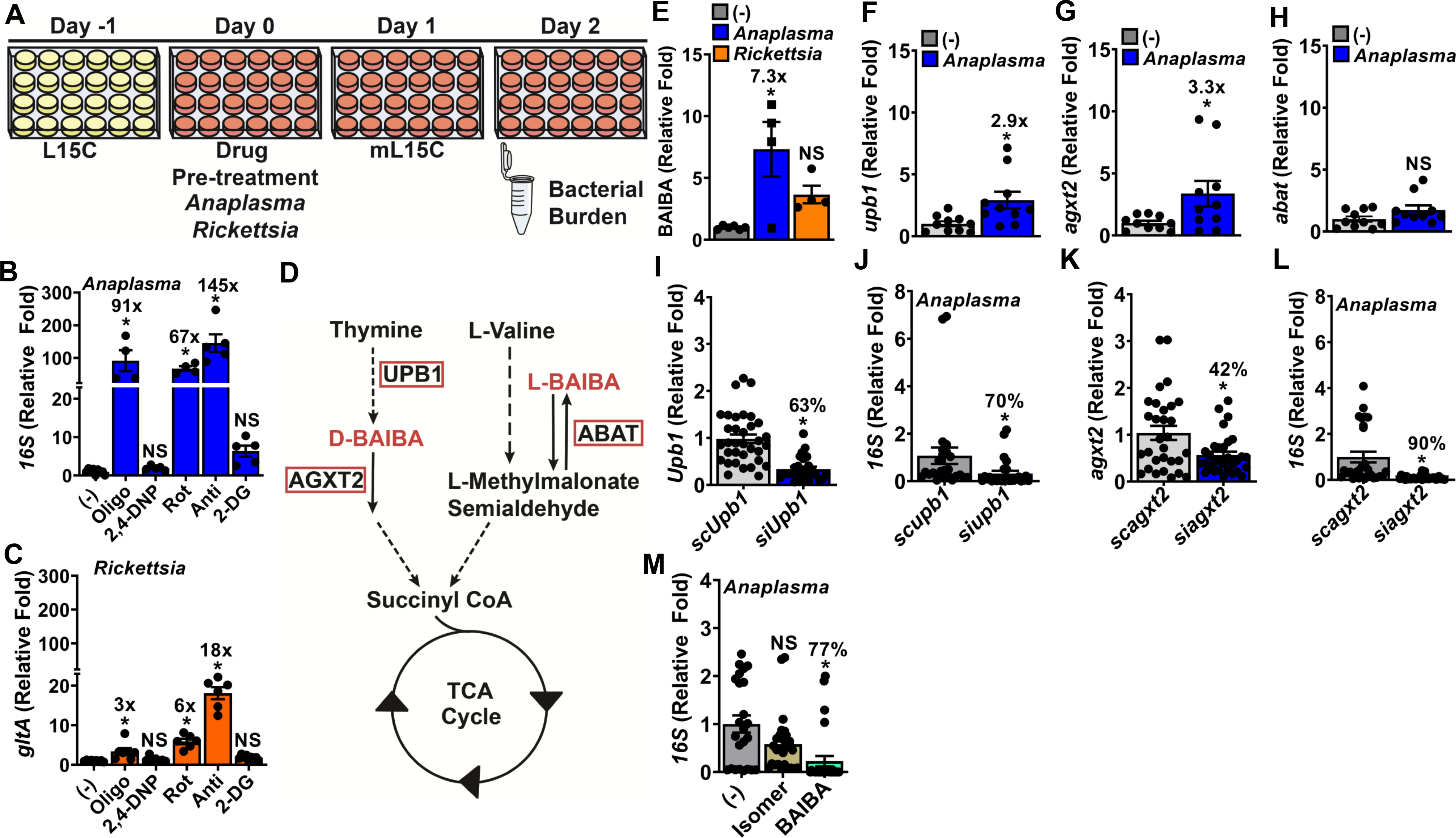
*A. phagocytophilum* depends on tick cell metabolites for *I. scapularis* infection. **A.** Schematics of microbial infection upon inhibitor treatment. **B**-**C** 1 x10^6^ ISE6 cells were treated with oligomycin – 0.5 µM; 2,4-DNP - 20 µM; rotenone – 0.1 µM; antimycin A – 0.5 µM or 2-DG – 50 mM 1 hour prior to infection. Cells were then incubated with (**B**) *A. phagocytophilum* (MOI 50) or (**C**) *R. buchneri* (MOI 50) for 48 hours. Data were normalized to untreated but infected control cells (-) to calculate fold changes in bacterial load. Data are representative of at least three independent experiments N=4-6. **D.** Schematics of BAIBA metabolism. Enzymes involved in catabolism and anabolism of BAIBA are highlighted in red. **E.** BAIBA levels measured in uninfected, *A. phagocytophilum*-infected, or *R. buchneri*-infected 5 x10^7^ ISE6 cells at MOI 50 24 hours post-infection N=4-6. **F-H.** Gene expression of ticks fed on uninfected mice or mice infected with *A. phagocytophilum*. Relative expression of (**F**) *upb1*, (**G**) *agxt2*, or (**H**) *abat* normalized to tick *actin*. Data are representative of two independent experiments N=10. **I-L.** Nymphs were injected with siRNA or scrambled control before feeding on *A. phagocytophilum*-infected mice for three days. Silencing efficiency in (**I**) *upb1* or (**K**) *agxt2* ticks. (**J** and **L**) *A. phagocytophilum* burden in silenced and control ticks. Bacterial burden was calculated by using the *A. phagocytophilum* specific *16S* rDNA gene and its relative expression normalized to *actin*. Data are representative of at least three independent experiments N=22-33. **M.** *A. phagocytophilum* burden in BAIBA-treated ticks. Nymphs were injected with 40 pmol of BAIBA, the α-aminoisobutyric acid (isomer) or phosphate-buffered saline (PBS) (-) and ticks were placed on *A. phagocytophilum*-infected mice for three days. Bacterial burden was calculated by using the *A. phagocytophilum* specific *16S* rDNA gene and its relative expression normalized to *actin*. Data are representative of two independent experiments N=24-28. Statistical significance was evaluated by one-way ANOVA followed by Dunnett’s post-hoc test (**B**, **C**, **E** and **M**) or unpaired t test with Welch’s correction (**F**-**L**). *, *p*<0.05. NS=not significant.

Given the results obtained for *A. phagocytophilum* and *R. buchneri* infection, we posited that these two obligate intracellular bacteria induced contrasting metabolic responses in *I. scapularis*. Hence, *I. scapularis* ISE6 cells were infected with either *A. phagocytophilum* or *R. buchneri* at indicated time points followed by an unbiased metabolomics analysis (Fig. S6) [data available via MetaboLights, identifier MTBLS686]. Metabolite levels were globally increased upon *A. phagocytophilum* infection, whereas *R. buchneri* contributed to fewer metabolic changes. Using a pathway enrichment analysis, we observed significant alterations in energy metabolites (figs. S7 and S8), nucleotide (figs. S9 and S10), fatty acid (figs. S11 and S12), methionine (figs. S13 and S14), protein degradation (figs. S15 and S16) and membrane lipid metabolism (fig. S17) upon *A. phagocytophilum* infection of *I. scapularis* ISE6 compared to *R. buchneri*. We concluded that *A. phagocytophilum* affects the metabolism of tick cells to a greater extent than the bacterial symbiont *R. buchneri*.

### D-β-aminoisobutyric acid (D-BAIBA) affects *A. phagocytophilum* acquisition in ticks

Next, we wanted to functionally characterize these pathways *in vivo* to demonstrate the utility of the metabolomics dataset for tick-microbe relationships. As described in the metabolomics analysis (figs. S6-S17), we determined that fatty acid, lipid and nucleotide metabolism in tick cells were impacted by *A. phagocytophilum* infection. Therefore, we focused our efforts on a pleiotropic metabolite that was involved in these processes. β-aminoisobutyric acid (BAIBA) is an intermediate of nucleotide and amino acid metabolism^44, 45^. BAIBA is also associated with fatty acid β-oxidation, lipid homeostasis and the browning of white adipose tissue in mammals^44, 45^. BAIBA has two enantiomers, D-BAIBA and L-BAIBA, that are generated by different enzymatic processes. During thymine degradation, N-carbamoyl BAIBA is converted to D-BAIBA by the enzyme β-ureidopropionase 1 (UPB1) before being catabolized to D- methylmalonate semialdehyde by alanine-glyoxylate aminotransferase 2 (AGXT2). Alternatively, L-BAIBA is a byproduct of L-valine catabolism and is generated by the enzyme 4-aminobutyrate aminotransferase (ABAT) through a reversible reaction from L-methylmalonate semialdehyde^45^. Both forms are eventually converted into propionyl-CoA, which funnels into the TCA cycle as a succinyl-CoA metabolite (Fig. 4D).

BAIBA was significantly elevated 24 hours after *A. phagocytophilum* infection compared to *R. buchneri* in *I. scapularis* ISE6 cells (Fig. 4E). Importantly, our metabolomics data did not distinguish between BAIBA enantiomers. Thus, we reconstructed the BAIBA pathway *in silico* and identified the ortholog genes in *I. scapularis* ticks for catabolism and anabolism: (*upb1*) [XM_029991952.1], (*agxt2*) [XM_029990918.1] and (*abat*) [XM_002405926.2] (Fig. 4D and table S2). We observed that ticks fed on *A. phagocytophilum*-infected mice upregulated the expression of *upb1* and *agxt2*, but not *abat* (Figs. 4F-H). Therefore, we characterized the impact of D-BAIBA regulation in tick-microbe interactions by manipulating the gene expression of *upb1* and *agxt2*. We silenced *upb1* or *agxt2* expression in ticks using small interfering RNAs (siRNAs) (Figs. 4I and 4K). RNAi remains the gold standard for disruption of tick proteins associated with biochemical pathways, as genome editing through clustered regularly interspaced short palindromic repeats (CRISPR) has only been applied to appendage genes to score morphological phenotypes^46^. *upb1-* or *agxt2*-silenced ticks fed on *A. phagocytophilum*-infected mice acquired significantly fewer bacteria than the control treatment (Figs. 4J and 4L). Then, we performed an experiment to determine whether the addition of exogenous BAIBA in ticks affects bacterial acquisition. We injected ticks with a racemic mixture of BAIBA and an isomer (α- aminoisobutyric acid) before placing these ectoparasites on mice infected with *A. phagocytophilum*. As shown in Figs. 4J and 4L, bacterial acquisition by BAIBA-injected ticks was significantly reduced compared to the isomer-injected ticks (Fig. 4M). Collectively, our findings indicated that D-BAIBA metabolism is important for *A. phagocytophilum* infection of ticks.

### Disruption of D-BAIBA-related enzymes affects tick fitness

Nucleotide metabolism is important for physiological and cellular homeostasis^47^. Given our observations with *A. phagocytophilum* colonization of ticks, we aimed to deconvolute the bacterial acquisition effect of D-BAIBA metabolism from tick fitness. Therefore, we silenced nymphs for *upb1* or *agxt2* (Figs. 5A and 5C) and measured attachment, feeding and survival. While we noticed a slight impairment in attachment for *I. scapularis* microinjected with the *upb1* siRNA (fig. S18), reduced weight in nymphs silenced for either *upb1* or *agxt2* was observed (Figs. 5B and 5D).

**Figure 5:**
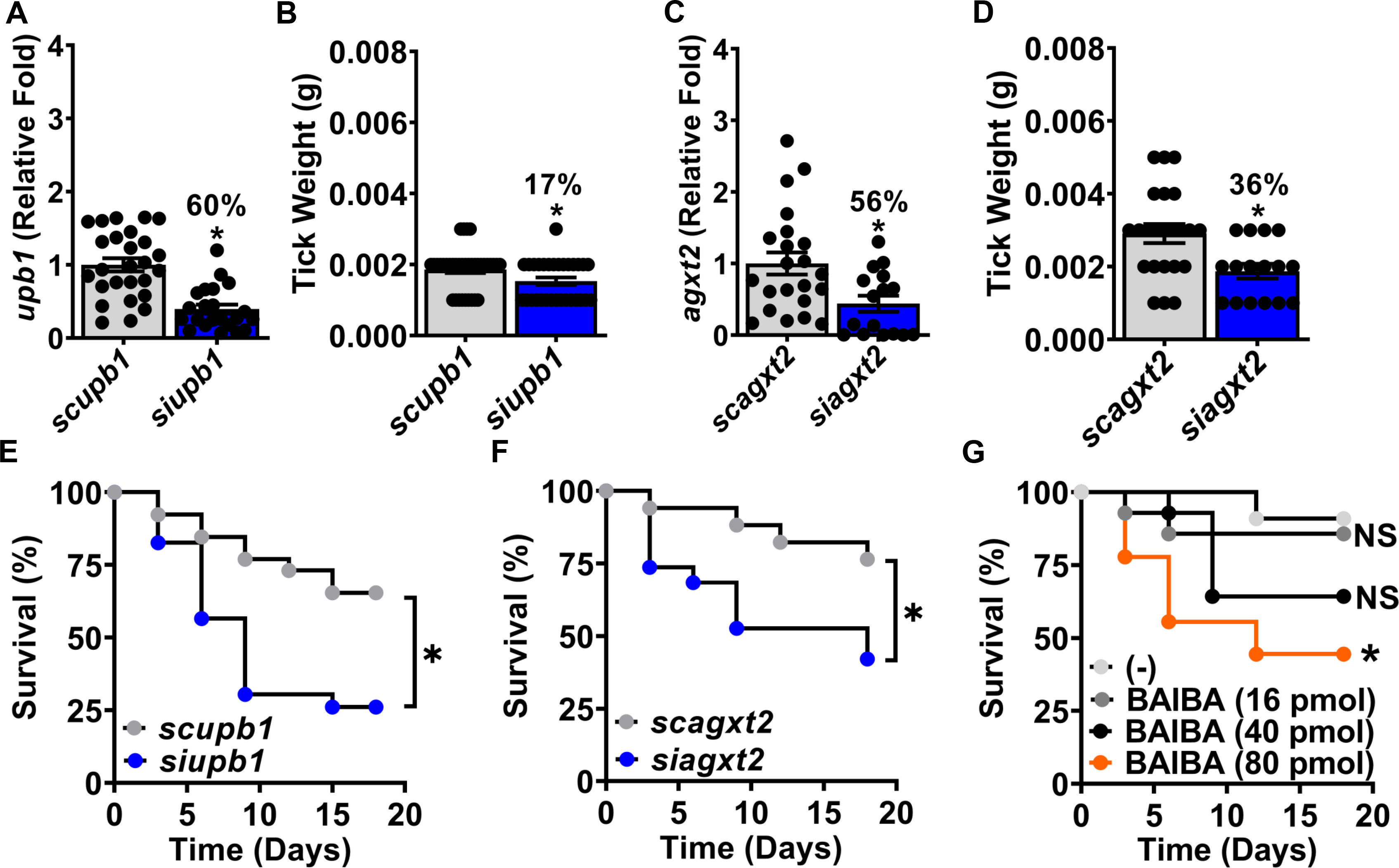
BAIBA regulates tick feeding and survival. A. Silencing efficiency of *upb1* in ticks. Nymphs were injected with the *upb1* siRNA (*siupb1*) or scrambled control (*scupb1*) before ticks were placed on uninfected mice to feed for three days. N=21-26. **B.** Weight of ticks post-feeding N=21-26. **C.** Silencing efficiency of *agxt2* in ticks. Nymphs were injected with the *agxt2* siRNA (*siagxt2*) or scrambled control sequence (*scagxt2*) before ticks were placed on uninfected mice to feed for three days. N=16-20. **D.** Weight of ticks post-feeding N=16-20. **E.** Survival of *siupb1-* or *scupb1*-injected ticks recorded 18 days post-blood meal N=23-26. **F.** Survival of *siagxt2-* or *scagxt2*-injected ticks recorded 18 days post-blood meal. N=17-19. Data are representative of at two independent experiments. **G.** Nymphs were injected with corresponding amounts of BAIBA before feeding on mice. Survival was recorded for 18 days. Data are representative of two independent experiments N=9-14. Statistical significance was evaluated by (**A-D**) unpaired t test with Welch’s correction or (**E-G**) Log rank (Mantel-Cox) test. *, *p*<0.05. NS=not significant.

Interestingly, the feeding deficiency was also detected when nymphs were microinjected with the *upb1* siRNA and fed on *A. phagocytophilum*-infected mice, but not in *agxt2*-silenced ticks (fig. S19). Nymphs silenced for *upb1* or *agxt2* exhibited reduced survival post-feeding compared to the control treatment (Figs. 5E-F). We then performed an experiment by microinjecting tick nymphs with exogenous amounts of racemic BAIBA to evaluate whether this metabolite influenced arthropod fitness. We used 16-80 pmols of racemic BAIBA in these experiments (fig. S20A-B) because 40 pmols successfully phenocopied the *siagxt2* siRNA injection treatment (Figs. 4K-M). There was no significant difference in attachment or weight for ticks injected with the racemic BAIBA when compared to the control treatment (fig. S20C-D). Conversely, we observed a dose-dependent decrease in survival for nymphs post-feeding (Fig. 5G).

Collectively, we determined that the enzymes involved in D-BAIBA metabolism affect tick feeding and survival. These results indicated a pleiotropic role for the tick metabolite BAIBA in tick fitness and bacterial acquisition.

## Discussion

In this study, we sought to understand how resource allocation and metabolism influence interspecies relationships. We developed a system for evaluating the metabolic status of tick cells and demonstrated how bioenergetic disruption affects infection. We determined that the ISE6 cell line is permissive to manipulation by small molecule inhibitors, which can be used for measuring glycolysis and cellular respiration in tick cells. Using an adapted metabolic flux assay, we observed that *A. phagocytophilum* and *B. burgdorferi* induce glycolysis but not OxPhos in tick cells. This phenomenon was distinct from *R. buchneri* endosymbiosis, suggesting a contrasting bacterial association in *I. scapularis*. In ticks, 2-DG and oligomycin exhibit detrimental effects on attachment and molting, respectively. These observations offer glimpses into the interconnection between bioenergetics and arthropod fitness, including how metabolic arrest impacts organismal homeostasis. The bioenergetics platform described in this study provides an invaluable tool for studying the metabolic interdependence between arthropod vectors and their microbial partners, a topic that is regrettably understudied.

Treatment with OxPhos inhibitors promoted a striking increase of bacterial burden in tick cells. Specifically, oligomycin, rotenone and antimycin A augmented *A. phagocytophilum* and *R. buchneri* load in *I. scapularis* ISE6 cells. It is plausible that arresting OxPhos promotes a buildup of substrates that rapidly accelerates bacterial division inside ticks. It is known that rickettsial bacteria carry small genomes and require host nutrients for survival^48–50^. Using a metabolomics approach, we compared changes that occur in tick cells infected with two obligate intracellular bacteria: *A. phagocytophilum* and *R. buchneri*. We found that *A. phagocytophilum* was more disruptive than the endosymbiont *R. buchneri*. It is possible that *A. phagocytophilum* draws more intracellular resources than *R. buchneri*; or, alternatively, it requires additional pathways for replication compared to *R. buchneri*. How ticks maintain cellular homeostasis when metabolites are elevated is not fully understood. Likewise, it remains to be determined if metabolite changes upon *A. phagocytophilum* infection are due to parasitism or a tick response to the microbe.

Ticks are hosts of a variety of symbionts, including species belonging to the genera *Coxiella, Rickettsia* and *Francisella*^31, 51^. Symbionts are essential for arthropod development and reproduction, as they provide essential vitamins and cofactors that ticks cannot sequester from imbibed blood (*e.g.*, biotin and folate)^51, 52^. *R. buchneri* encodes genes that may provide essential vitamins and cofactors and can be detected at high prevalence in *I. scapularis* populations^48, 53^, suggesting a possible selective advantage to the tick. This dependence indicates that tick symbionts have evolved mechanisms to ensure a balance of energy storage within the vector. This was noted in our metabolomics data, where we observed little to no alterations in tick energy metabolism during *R. buchneri* infection. In contrast, the disruption of tick metabolism by *A. phagocytophilum* is mirrored by its inability to be maintained transovarially among *Ixodes* spp. populations. Whether tick metabolism contributes to transstadial or transovarial transmission remains to be explored.

Given the metabolites uncovered in this study, we focused our efforts on hits that may be representative of several pathways. We discovered that silencing genes involved in the anabolism and catabolism of D-BAIBA reduced *A. phagocytophilum* burden and impaired tick feeding and survival. These findings indicate that D-BAIBA may be acting within a defined concentration window to enable tick fitness. We suggest that impairing D-BAIBA catabolism through the injection of *agxt2* siRNA in *I. scapularis* may lead to a metabolite hyper- accumulation and an alteration of the metabolic flux in ticks. On the other hand, altering D- BAIBA anabolism via the injection of *upb1* siRNA in *I. scapularis* may generate a hypo- accumulation of D-BAIBA. Decreased D-BAIBA levels in ticks likely inhibited downstream biochemical networks and homeostasis in *I. scapularis*. Altogether, these findings highlight the importance of a critical metabolite for life history programs in ticks, including maintenance, growth and survival^17–20^.

Finally, *A. phagocytophilum* altered metabolic pathways in ticks for efficient bacterial acquisition. In mammals, BAIBA is involved in thermoregulation by promoting the browning of adipose tissue^44^. Previous work demonstrated that *A. phagocytophilum* enhances tick survival through cold tolerance^13^. Whether BAIBA confers a survival advantage or contributes to thermoregulation of ticks remains unknown. Overall, we demonstrated that tools for studying metabolic activity in mammals can be applied to arthropod vectors and their microbial counterparts. Our findings highlight a shift in perspective for bioenergetics and resource allocation when evaluating microbial associations in arthropod vectors.

## Materials and Methods

### Cell culture

The *I. scapularis* IDE12 and ISE6, *A. americanum* AAE2 and *D. andersoni* DAE100 cell lines were obtained from Dr. Ulrike Munderloh at the University of Minnesota. All tick cell lines were maintained at 34°C in a non-CO2 incubator. Cells were cultured in T25 cm flasks (Greiner bio-one) containing L15C300 medium supplemented with 10% heat-inactivated fetal bovine serum (FBS, Millipore-Sigma), 10% tryptose phosphate broth (TPB, Difco), 0.1% bovine cholesterol lipoprotein concentrate (LPPC, MP Biomedicals). For *in vitro* experiments, ISE6 cells were plated in 48-well plates at a density of 1X10^6^ cells per well. The human leukemia cell line HL-60 was obtained from ATCC and maintained at 37°C in a 5% CO2 containing incubator.

Cells were cultured in T25 vented flasks (Cyto One) containing RPMI-1640 medium with L-Glutamine (Quality Biological) supplemented with 10% FBS (Gemini Bio-Products) and 1% GlutaMax (Gibco). All cell cultures were tested for *Mycoplasma* (Southern Biotech).

### Bacteria, mice and ticks

*A. phagocytophilum* strain HZ was grown as previously described in HL-60 cells at 37°C, using RPMI medium supplemented with 10% Fetal Bovine Serum and 1% Glutamax^54^. Bacterial numbers were calculated using the formula: number of infected HL-60 cells × 5 morulae/cell × 19 bacteria/cell × 0.5 (representing 50% recovery rate)^55^. Bacteria were purified by passing infected cells through a 27-gauge bent needle and using a series of centrifugation steps, as previously described^54^. Low passage isolate of *B. burgdorferi* B31 clone MSK5 was cultured in Barbour-Stoenner Kelly (BSK)-II medium supplemented with 6% normal rabbit serum at 34°C^56^, never exceeding 10^8^ bacteria per ml. Plasmid profiling was performed as described elsewhere^56^.

*R. buchneri* strain ISO7^T^ was obtained from Dr. Ulrike Munderloh. *R. buchneri* was maintained in ISE6 cells at 30°C in a non-CO2 incubator^6^. Bacteria were isolated from infected ISE6 cells using a 27-gauge needle and cell debris was separated by centrifugation at 600xg for 10 mins. Spirochetes were counted using a light- or dark-field (Zeiss Primo Star Microscope) under a 40X objective lens, respectively^56^.

Age matched, six- to ten-week-old C57BL/6J male mice were supplied by the University of Maryland Veterinary Resources or Jackson Laboratories. *I. scapularis* nymphs were obtained from either Oklahoma State University or the University of Minnesota breeding colonies. Upon arrival, ticks were housed in an incubator at 23°C with >85% relative humidity and a 14/10-hour light/dark photoperiod regimen. Animal experiments were approved by the Institutional Biosafety (IBC, IBC-00002247) and Animal Care and Use (IACUC, #0119012) committees at the University of Maryland School of Medicine and complied with National Institutes of Health (NIH) guidelines (Office of Laboratory Animal Welfare [OLAW] assurance number A3200-01).

### RNA interference and tick injection experiments

Small interfering RNAs (siRNA) and their scrambled controls (scRNA) were synthesized using the Silencer siRNA construction kit (Thermo Scientific) according to the manufacturer’s instructions. Nymphs were microinjected with 30-50 ng of siRNA or scRNA, as previously described^57^. Ticks were allowed to recover overnight before being placed on uninfected or *A. phagocytophilum-*infected C57BL/6J mice. Ticks were collected 3 days after placement in which the degree of host attachment and tick weight were assessed. Ticks were either placed in a incubator (23^°^C, with >85% relative humidity in a 14/10-hour light/dark photoperiod regimen) for survival experiments lasting 18 days or frozen at -80°C in 200 ml TRIzol reagent for RNA extraction.

For BAIBA and inhibitor treatments, ticks were microinjected with 60-80 nl of BAIBA or inhibitor solution and allowed to recover overnight. The following day, ticks were placed on anesthetized mice. Three days after placement, tick attachment and weight were recorded, and collected ticks were either placed in a humidified chamber for survival/molting experiments or frozen in TRIzol reagent for RNA extraction. For molting experiments, *I. scapularis* were monitored until ticks in the control treatment molted.

### Mouse infections

Age matched, six- to ten-week-old C57BL/6J male mice were used for *A. phagocytophilum* acquisition experiments. *A. phagocytophilum* was isolated from infected HL-60 cells and resuspended in PBS at a concentration of 1x10^8^ bacteria per ml. Mice were intraperitoneally injected with 100 μl of the inoculum (1×10^7^ total *A. phagocytophilum*). Infection progressed for 7 days before placing ticks.

### Bioenergetic measurements in ISE6 cells using the Seahorse analyzer

OCR and ECAR were measured in XF96 cell culture microplates using a Seahorse XFe96 Extracellular Flux Analyzer (Seahorse, Agilent Technologies). ISE6 cells were seeded at densities of 120 – 150,000 cells per well in complete L15C300 media^41^ and incubated at 34°C for 24 hours. Media was then replaced with modified L15C media (mL15C) alone or mL15C containing *A. phagocytophilum*, *B. burgdorferi* or *R. buchneri* for 48 hours (table S1). For OCR detection, values were measured at basal conditions and after 20 µM 2,4-DNP (Sigma Aldrich), 0.1 µM rotenone (Sigma Aldrich) and 0.5 µM antimycin A (Sigma Aldrich) treatments. For ECAR detection, values were measured at basal conditions and after adding 50 mM glucose and 2-DG (Sigma Aldrich). Data normalization was performed using a Celigo image cytometer (Nexcelom Bioscience, Massachusetts) following the manufacturer guidelines for measuring cell confluency. The cartridge was calibrated with the Seahorse XF Calibrant Solution (Agilent Technologies) at 37°C in a non-CO2 and non-humidified incubator for at least 2 h prior to the assay.

### qRT-PCR analysis

Tick and cell samples were preserved in the TRIzol reagent prior to RNA extraction.

Total RNA was isolated using the PureLink RNA Mini kit (Ambion). cDNA was synthesized from 300-600 ng RNA using the Verso cDNA Synthesis kit (ThermoFisher). Gene expression was measured using a CFX96 Touch Real-Time PCR Detection System (Bio-rad) with iTaq Universal SYBR Green Supermix (Bio-rad). Expression levels for genes were calculated by relative quantification normalized to tick *actin*. Primers used are listed in table S2.

### Metabolomics

To generate samples for metabolomics, 5x10^7^ ISE6 cells were placed in T25 flasks in L15C300 media. The following day, media was removed and replaced with mL15C media alone or mL15C media containing either *A. phagocytophilum* or *R. buchneri* at a MOI 50. Cells were harvested via a cell scraper at 1 hour and 24 hours post-infection followed by centrifugation at 3,320xg for 10 minutes at 4°C. Cell pellets were frozen in liquid nitrogen and shipped to Metabolon Inc. for analysis. Four independent experiments were performed for each condition, and data were normalized according to protein concentrations.

Sample processing was performed at Metabolon Inc., as previously described^58^.

Individual samples were subjected to methanol extraction and then separated into aliquots for ultra-high performance liquid chromatography/mass spectrometry (UHPLC/MS). Global biochemical profiling involved reverse phase chromatography positive ionization for hydrophilic (LC/MS Positive Polar) and hydrophobic (LC/MS Positive Lipid) compounds, reverse phase chromatography with negative ionization (LC/MS Negative), and a hydrophilic interaction chromatography (HILIC) coupled to negative ion mode electrospray ionization (LC/MS Polar)^59^. Methods interspersed between full mass spectrometry and refragmentation (MSn) scans.

Metabolites were identified by automated comparison of the ion features in the experimental samples to a reference library of at least 4000 chemical standard entries^60^.

Metabolon Inc. performed the initial statistical analysis for the metabolite study. Two types of statistical analyses were performed: (1) significance tests and (2) classification analysis. Standard statistical analyses were performed in ArrayStudio on log-transformed data. For non-standard analyses RStudio was used. Following log transformation, Welch’s two sample *t*-test identified biochemicals that differed significantly (*p* < 0.05) and false discovery rate (*q* value) were calculated between treatments. Principal components analysis, hierarchical clustering, and random forest were used for metabolite classification. Time points were equaled to 1 and each compound in the original scale (raw area count) was rescaled to set the median across samples. Data are available via MetaboLights, identifier MTBLS686.

### Colorimetric assays

ISE6 cells were seeded in 48-well microtiter plate (Cyto-one) with complete L15C300 medium at a density of 1x10^6^ for 24 hours. Cells were challenged with *A. phagocytophilum*, *B. burgdorferi* or *R. buchneri* (MOI 50) in mL15C media supplemented with 10 mM glucose and incubated for 1 or 24 hours. Cells were harvested and resuspended in assay buffer. Glycolytic activity was evaluated using kits measuring lactate (Sigma Aldrich), lactate dehydrogenase (LDH, Sigma Aldrich), phosphoglucose isomerase (PGI, Sigma Aldrich) and nicotinamide adenine dinucleotide (NAD/NADH, Sigma Aldrich). TCA cycle activity was evaluated from kits measuring citrate (Sigma Aldrich), aconitase activity (Sigma Aldrich), succinate (Sigma Aldrich) and succinate dehydrogenase activity (SDH, Sigma Aldrich). Reagents are listed in table S3.

### Statistical analysis

Statistical significance between two conditions were assessed using an unpaired *t*-test with Welch’s correction for unequal variances. One-way ANOVA followed by the Dunnett’s multiple comparisons test was used for analyzing statistical differences between three or more groups. For categorical variables, Fisher’s Exact or Chi-square test was used. Survival curves were analyzed with the Log-rank (Mantel-Cox) test. All statistical analysis were performed in GraphPad PRISM® (GraphPad Software version 9.1.0). Outliers were detected by a Graphpad Quickcals program (https://www.graphpad.com/quickcalcs/Grubbs1.cfm).

## Acknowledgements

We thank Ulrike G. Munderloh (University of Minnesota) for providing tick cell lines and Jonathan Oliver (University of Minnesota) for providing *I. scapularis* nymphs; Joseph Gillespie (University of Maryland, Baltimore) and Timothy Driscoll (West Virginia University) for sharing primer details to measure *R. buchneri*; Dana Shaw (Washington State University) and Adela Oliva Chavez (Texas A&M University) for information regarding cultures of *B. burgdorferi* and *A. phagocytophilum*, respectively; Erin E. McClure Carroll (University of Maryland School of Medicine) for schematics and Holly Hammond for administrative support; and the Biopolymer/Genomics core facility for Sanger sequencing and the Seahorse metabolic flux assay (S10OD025101). This work was supported by grants from the National Institutes of Health (NIH) to AJO (F31AI152215), LRB (F31AI167471), HJL (T32AI162579), JHFP (R01AI134696, R01AI116523, R01AI049424 and P01AI138949). JHFP was also supported in-kind by the Fairbairn Family Lyme Research Initiative. The content is solely the responsibility of the authors and does not necessarily represent the official views of the NIH, the Department of Health and Human Services, or the United States government.

## Author contributions

SS and JHFP designed the study. SS, AR, LM, AJO, NS, XW, HJL, LRB, FECP and PR performed the experiments. SS, AJO, HJL, and JHFP wrote the manuscript. LRB aided with experimentation and created some schematics. All authors analyzed the data, provided intellectual input into the study, and contributed to editing of the manuscript. GMF and BMP supervised experiments and provided instruments. JHFP supervised the study.

