## Supplementary Information for "Metabolic disruption impacts tick fitness and microbial relationships"

**This file includes:**

Figures S1-S20

Tables S1-S3

#### Supplementary Figure Legends

**Figure S1: Viability of tick ISE6 cells in cultured media.** $1 \times 10^6$ cells cultured in L15C300 (L15C) complete or modified L15C (mL15C) media for 48 hours. Statistical significance was evaluated by the unpaired t test with Welch's correction. NS = not significant. Data are representative of at least two independent experiments.

**Figure S2: Drug susceptibility of tick cell lines.** Viability of (**A-E**) *I. scapularis* (IDE12), (**F-J**) *A. americanum* (AAE2) and (**K-O**) *D. andersoni* (DAE100) cell lines treated with different inhibitor concentrations (N=4). Blue bars indicate non-significant, whereas red bars denote a statistically significant decrease in cell viability. Data are representative of two independent experiments. Statistical significance was calculated by one-way ANOVA followed by Dunnett's test using multiple comparisons. $p<0.05$.

**Figure S3: 2-DG does not affect tick survival, feeding, or molting.** (**A**) Survival curve of unfed ticks treated with 2-DG. Data are representative of two independent experiments. (**B**) Survival curve of fed ticks treated with 2-DG. Ticks were injected with 2-DG prior to feeding for three days. Data are representative of two independent experiments. (**C**) Weight of fed ticks treated with 2-DG. Ticks were injected with 2-DG prior to feeding for three days. (**D**) Molting of ticks treated with 2-DG. Ticks were injected with 2-DG prior to feeding for three days. (**E**) Percentage of ticks that molted in (**D**). Data are representative of two independent experiments. Statistical significance was evaluated by (**A**, **B**, **D**) Log-rank (Mantel-Cox) test, (**C**) One-way ANOVA followed by Dunnett's test with multiple comparisons or (**E**) Chi-square test. $p<0.05$. NS=not significant.

**Figure S4: Oligomycin injection in the tick *I. scapularis*.** (**A**) Survival curve of unfed ticks injected with oligomycin. (**B**) Percentage of ticks that attached after oligomycin injection. Ticks were injected with 0.8 pmol oligomycin and placed on mice overnight. (**C**) Survival curve of ticks treated with oligomycin. Ticks were injected with 0.8 pmol oligomycin prior to feeding for three days. (**D**) Weight of ticks treated with oligomycin. Ticks were injected with 0.8 pmol oligomycin prior to feeding for three days. Data are representative of two independent experiments. Statistical significance was evaluated by (**A, C**) Log-rank (Mantel-Cox), (**B**) Fisher's exact test or (**D**) the unpaired t test. *, $p<0.05$. NS=not significant.

**Figure S5: *A. phagocytophilum* infection does not alter the tick TCA cycle.** (**A**) Abbreviated representation of glycolysis (green), TCA (pink) and OxPhos (blue) with measured TCA components highlighted in red. (**B-E**) $1 \times 10^6$ ISE6 cells were stimulated with *A. phagocytophilum* (MOI 50) or left untreated for 24 hours. (**B**) Citrate, (**C**) aconitase activity, (**D**) succinate and (**E**) succinate dehydrogenase (SDH) activity were measured by colorimetric assays. N=5. Data are representative of two independent experiments. Statistical significance was evaluated by the unpaired t test. $p<0.05$. NS=not significant.

**Figure S6: Schematics of the unbiased metabolomics analysis in tick cells.** *I. scapularis* ISE6 ($5 \times 10^7$) cells were stimulated for 1 or 24 hours with either *A. phagocytophilum* or *R. buchneri* at multiplicity of infection (MOI) 50. Mass spectrometry was followed by metabolite identification and pathway analysis, as described in the materials and methods [data available via MetaboLights, identifier MTBLS686].

**Figure S7: Schematics of glycolysis and the TCA cycle pathway.**

**Figure S8: Measurement of bioenergetics in tick cells after microbial infection.** Tick ISE6 cells were untreated (grey), infected with *A. phagocytophilum* (MOI 50) or the endosymbiont *R. buchneri* (MOI 50) for 1 and 24 hours. N=4-6. Statistical significance was evaluated by one-way ANOVA followed by the Dunnett's test for multiple comparisons. $p<0.05$. NS=not significant.

**Figure S9: Schematics of nucleotide metabolism pathways.**

**Figure S10: Nucleotide metabolism in tick cells after microbial infection.** Relative metabolite levels associated with nucleotide metabolism after microbial infection. ISE6 cells were left untreated (grey), infected with *A. phagocytophilum* (MOI 50) or the endosymbiont *R. buchneri* (MOI 50) for 1 (left side) or 24 (right side) hours. N=4-6. Statistical significance was evaluated by one-way ANOVA followed by the Dunnett's test for multiple comparisons. *, $p<0.05$. NS=not significant.

**Figure S11: Schematics of fatty acid metabolism.**

**Figure S12: Fatty acid metabolism of tick cells after microbial infection.** Relative metabolite levels associated with fatty acid metabolism after microbial infection. ISE6 cells were left untreated (grey), infected with *A. phagocytophilum* (MOI 50) or the endosymbiont *R. buchneri* (MOI 50) for 1 (left side) and 24 (right side) hours. N=4-6. Statistical significance was evaluated by one-way ANOVA followed by Dunnett's test for multiple comparisons. *, $p<0.05$. NS=not significant.

**Figure S13: Schematics of methionine metabolism.**

**Figure S14: Methionine metabolism of tick cells after microbial infection.** Relative metabolite levels associated with methionine metabolism after microbial infection. ISE6 cells were left untreated (grey), infected with *A. phagocytophilum* (MOI 50) or the endosymbiont *R. buchneri* (MOI 50) for 1 (left side) and 24 (right side) hours. N=4-6. Statistical significance was evaluated by one-way ANOVA followed by Dunnett's test for multiple comparisons. *, $p<0.05$. NS=not significant.

**Figure S15: Schematics of protein degradation products.**

**Figure S16: Protein degradation in tick cells after microbial infection.** Relative metabolite levels associated with protein degradation after microbial infection. ISE6 cells were left untreated (grey), infected with *A. phagocytophilum* (MOI 50) or the endosymbiont *R. buchneri* (MOI 50) for 1 (left side) and 24 (right side) hours. N=4-6. Statistical significance was evaluated by one-way ANOVA followed by Dunnett's test for multiple comparisons. *, $p<0.05$. NS=not significant.

**Figure S17: Membrane lipid metabolism in tick cells after microbial infection.** Relative metabolite levels associated with membrane lipid metabolism after microbial infection. ISE6 cells were left untreated (grey), infected with *A. phagocytophilum* (MOI 50) or the endosymbiont *R. buchneri* (MOI 50) for 1 (left side) and 24 (right side) hours. N=4-6. Statistical significance was evaluated by one-way ANOVA followed by Dunnett's test for multiple comparisons. *, $p<0.05$. NS=not significant.

**Figure S18: Tick attachment after silencing of genes associated with D-BAIBA metabolism.** (**A**) *I. scapularis* nymphs were injected with *upb1* siRNA (*siupb1*) or the scrambled control sequence (*scupb1*) and placed on uninfected mice overnight. N=21-26. Attachment was measured. (**B**) *I. scapularis* nymphs were injected with *agxt2* siRNA (*siagxt2*) or scrambled control sequence (*scagxt2*) siRNA and placed on uninfected mice overnight. Attachment was recorded N=16-20. Statistical significance was evaluated by the Fisher's exact test. *, $p<0.05$. NS=not significant. Data are representative of two independent experiments.

**Figure S19: Fitness in ticks silenced for D-BAIBA metabolism genes and fed on *A. phagocytophilum*-infected mice.** (**A**) Attachment and (**B**) weight of *I. scapularis* nymphs injected with *upb1* siRNA (*siupb1*) or the scrambled control sequence (*scupb1*) and placed on *A. phagocytophilum*-infected mice for three days N=22-33. (**C**) Attachment and (**D**) weight of *I. scapularis* nymphs injected with *agxt2* siRNA (*siagxt2*) or scrambled control sequence (*scagxt2*) and placed on *A. phagocytophilum*-infected mice for three days N=35-38**.** Statistical significance was evaluated by the (**A, C**) Fisher's exact test or the (**B, D**) unpaired t test with Welch's correction. *, $p<0.05$. NS=not significant. Data are representative of two independent experiments.

**Figure S20: Exogenous BAIBA injection has no effect on tick attachment and feeding.** (**A**) *I. scapularis* nymphs were injected with corresponding amounts of BAIBA and placed on mice for three days. Tick (**A**) attachment and (**B**) weight were recorded N=9-15. (**C** and **D**) Nymphs were injected with phosphate buffered saline (-), an isomer of BAIBA or BAIBA itself (40 pmol each) and placed on mice for three days. (**C**) Attachment and (**D**) weight of ticks were recorded N=21-28. Data represents at least two independent experiments. Statistical significance was evaluated by the (**A** and **C**) Chi-square test or (**B** and **D**) One-way ANOVA followed by Dunnett's test for multiple comparisons. $p<0.05$. NS=not significant.

**Table S1. Composition of the modified tick cell medium for the Seahorse assay**

| Seahorse DMEM | Modified L15C (mL15C) | L15C300 Complete |
|---|---|---|
| No glucose | **No glucose** | 100mM glucose |
| 2 mM glutamine | **2 mM glutamine** | 5.4mM glutamine |
| 1 mM sodium pyruvate | *5 mM sodium pyruvate | 5mM sodium pyruvate |
| No D-galactose | *5 mM D-galactose | 5mM D-galactose |
| No bicarbonate | No bicarbonate | No bicarbonate |
| No α-ketoglutarate | *3.0 mM α-ketoglutarate | 3.0mM α-ketoglutarate |
| No glutamic acid | *3.3 mM glutamic acid | 3.3mM glutamic acid |
| No proline | *3.9 mM proline | 3.9mM proline |
| No fetal bovine serum (FBS) | **No fetal bovine serum (FBS)** | 10 % fetal bovine serum (FBS) |
| No tryptose phosphate broth (TPB) | **No tryptose phosphate broth (TPB)** | 10 % tryptose phosphate broth (TPB) |
| No lipoprotein-cholesterol concentrate (LPPC) | **No lipoprotein-cholesterol concentrate (LPPC)** | 0.01 % lipoprotein-cholesterol concentrate (LPPC) |
| pH = 7.4 | **pH = 7.4** | pH = 6.0-7.0 |

A modified medium was developed for studying the metabolic flux in tick cells through the Seahorse analyzer. *Components included in the L15C300 complete medium compared to the standard DMEM medium for the Seahorse assay in mammalian cells. Bold denotes the difference in chemical composition between the mL15C and the L15C300 complete medium used to culture tick cells.

**Table S2: Primers and siRNA sequences**

| Target | Type | Start position Within mRNA | siRNA/Primer Name | Strand | Primer sequence | Accession number |
|---|---|---|---|---|---|---|
| *I. scapularis upb1** | siRNA_1 (combination of siRNA_1,2 and 3 was used for all *in vitro* and *in vivo* experiments) | 253 | *siupb1*_253F | forward | AAGAAATCCAGAAAGTGGTCGCCTGTCTC | XM_029991952.1 |
| | | | *siupb1*_253R | reverse | AACGACCACTTTCTGGATTTCCCTGTCTC | |
| | siRNA_2 | 430 | *siupb1*_430F | forward | AAGATTCTCAAGGAGCACCTTCCTGTCTC | |
| | | | *siupb1*_430R | reverse | AAAAGGTGCTCCTTGAGAATCCCTGTCTC | |
| | siRNA_3 | 735 | *siupb1*_735F | forward | AACATCATCTGCTTTCAGGAGCCTGTCTC | |
| | | | *siupb1*_735R | reverse | AACTCCTGAAAGCAGATGATGCCTGTCTC | |
| | scramble_1 (combination of scramble_1,2 and 3 was used for all *in vitro* and *in vivo* experiments) | | *scupb1*_253F | forward | AAGGCAGTGCGAATAAAGACTCCTGTCTC | |
| | | | *scupb1*_253R | reverse | AAAGTCTTTATTCGCACTGCCCCTGTCTC | |
| | scramble_2 | | *scupb1*_430F | forward | AAGCTGACGGTCGGCACACCTCCTGTCTC | |
| | | | *scupb1*_430R | reverse | AAAGGTGTGCCGACCGTCAGCCCTGTCTC | |
| | scramble_3 | | *scupb1*_735F | forward | AAGAGCCTCACTCTTAGTTGACCTGTCTC | |
| | | | *scupb1*_735R | reverse | AATCAACTAAGAGTGAGGCTCCCTGTCTC | |
| | qRT-PCR | 608 | | forward | GAGTGGTGCGTGTCGGGCTTGTGC | |
| | | 687 | | reverse | CACCACACAGGGCAGCGGTGTCC | |
| *I. scapularis agxt2** | scramble_1 (combination of scramble_1,2 and 3 was used for all *in vitro* and *in vivo* experiments) | 202 | *siagxt2*_202F | forward | AAGAACATGTCCAGGTATCGCCCTGTCTC | XM_029990918.1 |
| | | | *siagxt2*_202R | reverse | AAGCGATACCTGGACATGTTCCCTGTCTC | |

| | | | | | | |
|---|---|---|---|---|---|---|
| | siRNA_2 | 364 | *siagxt2*_364F | forward | AACTTGAGATTTCCGGGCAACCCTGTCTC | |
| | | | *siagxt2*_364R | reverse | AAGTTGCCCGGAAATCTCAAGCCTGTCTC | |
| | siRNA_3 | 952 | *siagxt2*_952F | forward | AAAAAGCCGTTGCCGATTCCTCCTGTCTC | |
| | | | *siagxt2*_952R | reverse | AAAGGAATCGGCAACGGCTTTCCTGTCTC | |
| | scramble_1 (combination of scramble_1,2 and 3 was used for all *in vitro* and *in vivo* experiments) | | *scagxt2*_202F | forward | AAACCTATGGTATACGCGGACCCTGTCTC | |
| | | | *scagxt2*_202R | reverse | AAGTCCGCGTATACCATAGGTCCTGTCTC | |
| | scramble_2 | | *scagxt2*_364F | forward | AATAGATCCTGTAACCGTGGCCCTGTCTC | |
| | | | *scagxt2*_364R | reverse | AAGCCACGGTTACAGGATCTACCTGTCTC | |
| | scramble_3 | | *scagxt2*_952F | forward | AAAGTACGTCCCGATTTGCACCCTGTCTC | |
| | | | *scagxt2*_952R | reverse | AAGTGCAAATCGGGACGTACTCCTGTCTC | |
| | qRT-PCR | 324 | | forward | GTATCCATCGGTCCACGAGT | |
| | | 420 | | reverse | CCAGCATCATGGCTAAGTCG | |
| *I. scapularis abat** | siRNA_1 (combination of siRNA_1,2 and 3 was used for all *in vitro* and *in vivo* experiments) | 1293 | *siabat*_1293F | forward | AAATCAGAGGCCCTGAAGAGCCCTGTCTC | XM_002405926.2 |
| | | | *siabat*_1293R | reverse | AAGCTCTTCAGGGCCTCTGATCCTGTCTC | |
| | siRNA_2 | 1650 | *siabat*_1650F | forward | AAGGCCGTGTTCATCAACTACCCTGTCTC | |
| | | | *siabat*_1650R | reverse | AAGTAGTTGATGAACACGGCCCCTGTCTC | |
| | siRNA_3 | 1824 | *siabat*_1824F | forward | AAGGCCATCCATAAACTGGACCCTGTCTC | |
| | | | *siabat*_1824R | reverse | AAGTCCAGTTTATGGATGGCCCCTGTCTC | |
| | scramble_1 (combination of scramble_1,2 and 3 was used for all *in vitro* and *in vivo* experiments) | | *scabat*_1293F | forward | AAGCCAACCGGCTGAGTGAAACCTGTCTC | |
| | | | *scabat*_1293R | reverse | AATTTCACTCAGCCGGTTGGCCCTGTCTC | |
| | scramble_2 | | *scabat*_1650F | forward | AAGACGGACGTACCATAGGTTCCTGTCTC | |

| | | | | | | |
|---|---|---|---|---|---|---|
| | | | *scabat*_1650R | reverse | AAAACCTATGGTACGTCCGTCCCTGTCTC | |
| | scramble_3 | | *scabat*_1824F | forward | AAGCCACAACTAGGGACATCTCCTGTCTC | |
| | | | *scabat*_1824R | reverse | AAAGATGTCCCTAGTTGTGGCCCTGTCTC | |
| | qRT-PCR | 2006 | | forward | GGTCGAGCCAATCCAGGCCGAAG | |
| | | 2107 | | reverse | CACAGCCGGTCTGGACTTCGTCGC | |
| *I. scapularis β-actin*[#] | qRT-PCR | 896 | *I.s. actin*_F | forward | GGTATCGTGCTCGACTC | XM_029977298 |
| | | 1003 | *I.s. actin*_R | reverse | ATCAGGTAGTCGGTCAGG | |
| *A. phagocytophilum 16S*[†] | qRT-PCR | 116 | *Ap 16S*_F | forward | GGTGAGTAATGCATAGGAATC | APH_RS03965# |
| | | 223 | *Ap 16S*_R | reverse | GCTCATCTAATAGCGATAAATC | |
| *R. buchneri gltA*[¥] | qRT-PCR | 1127 | *Rb gltA*_F | forward | TCGCAAATGTTCACGGTACTTT | CP006009.1 |
| | | 1179 | *Rb gltA*_R | reverse | TCGTGCATTTCTTTCCATTGTG | |

*siRNA designed prior to the recently published *I. scapularis* reference genome (De *et al.*, 2023 doi: 10.1038/s41588-022-01275-w).

[#]Shaw et al., 2017; DOI:10.1038/ncomms14401

[†]Oliva Chavez et al., 2021; DOI:10.1038/s41467-021-23900-8

[¥]Stenos et al., 2005; DOI:10.4269/ajtmh.2005.73.1083

**Table S3: Resources and reagents available**

| Cell Medium | Source | Identifier | Dilution/Concentration |
|---|---|---|---|
| Leibovitz's L-15 Medium, Powder | Gibco | 41300039 | |
| Fetal Bovine Serum (for tick cells) | Millipore-Sigma | F0926-500ML | 10% |
| Tryptose Phosphate Broth | Difco | 260300 | 10% |
| Bovine Cholesterol Lipoprotein Concentrate | MP Biomedicals | 191476 | 0.1% |
| RPMI-1640 Medium With L-Glutamine | Quality Biological | 112-025-101 | |
| Fetal Bovine Serum (for HL-60s) | Gemini Bio-Products | 100-106 | 10% |
| GlutaMax | Gibco | 35050-061 | 1% |
| CMRL1066 w/L-Glutamine (Powder) | US Biological | C5900 | |
| Sodium Citrate Tribasic Dihydrate | Millipore-Sigma | S4641 | 0.7g/L |
| Yeastolate | BD | 255772 | 2.0g/L |
| Neopeptone | BD | 211681 | 5.0g/L |
| N-Acetyl-α-D-glucosamine | Millipore-Sigma | 1079-25GM | |
| Albumin, Bovine Fraction V | MP Biomedicals | 160069 | |
| Sodium Pyruvate | Millipore-Sigma | P5280 | 0.8 g/L |
| Distilled Water | Gibco | 15-230-147 | |
| L-aspartic Acid | Millipore-Sigma | 11189 | 0.449 g/L |
| L-glutamine | Millipore-Sigma | G8540 | 0.500 g/L |
| L-proline | Millipore-Sigma | 81709 | 0.450 g/L |
| L-glutamic Acid | Millipore-Sigma | 49449 | 0.250 g/L |
| Alpha-ketoglutaric Acid | Millipore-Sigma | K1128 | 0.449 g/L |
| Sodium Hydroxide | Millipore-Sigma | S8045 | 10N (stock) |
| **Materials** | | | |
| Seahorse Fluxpaks | Agilent Technologies | 102416-100 | |
| Miller GP 0.2 µm Filter Unit | Millipore-Sigma | SLGP033RS | |
| 1.5 ml Microcentrifuge Tubes | Thomas Scientific | 1148T71 | |
| 15 ml Conical Screw Cap Tubes | USA Scientific | 5618-8261 | |
| 50 ml Conical Screw Cap Tubes | USA Scientific | 5622-7270 | |
| 500 ml Vacuum Filter/Storage Bottle, 0.2 µm | Corning | 430773 | |
| 27 Gauge, 1/2" Needle | BD | 305109 | |
| T-25 Flasks (for tick cells) | Greiner bio-one | 690160 | |
| T-25 Vented Flasks (for HL-60s) | Cyto One | CC7682-4825 | |

| | | | |
|---|---|---|---|
| Nonstick Microcentrifuge Tubes | Ambion | AM12450 | |
| Microscope Slides | Fisher Scientific | 12550400 | |
| 48-well Microtiter Plate | Cyto-one | CC7682-7548 | |
| **Reagents** | | | |
| Ethyl alcohol, Pure; 200 proof for molecular biology | Millipore-Sigma | E7023-1L | 70-100% |
| TRIzol Reagent | Ambion | 15596018 | |
| iTaq Universal SYBR Green Supermix | Bio-rad | 1725121 | |
| HyClone™ Water, Molecular Biology Grade | Cytiva | SH3053801 | |
| Oligomycin | Sigma Aldrich | 75351 | 0.5μM |
| 2,4-dinitrophenol | Sigma Aldrich | D198501 | 20μM |
| Rotenone | Sigma Aldrich | 45656 | 0.1μM |
| Antimycin A | Sigma Aldrich | A8674 | 0.5μM |
| 2-deoxy-D-glucose | Sigma Aldrich | D6134-5G | 50mM |
| **Commercial Assays** | | | |
| Mycoplasma Test Kit | Southern Biotech | 13100-01 | |
| Silencer siRNA Construction Kit | Thermo Scientific | AM1620 | |
| PureLink RNA Mini Kit | Ambion | 12183025 | |
| Verso cDNA Synthesis Kit | ThermoFisher | AB-1453 | |
| Lactate Assay Kit | Sigma Aldrich | MAK064 | |
| Lactate Dehydrogenase Activity Assay Kit | Sigma Aldrich | MAK066 | |
| Phosphoglucose Isomerase Colorimetric Assay Kit | Sigma Aldrich | MAK103 | |
| NAD/NADH Quantification Kit | Sigma Aldrich | MAK037 | |
| Citrate Assay Kit | Sigma Aldrich | MAK057 | |
| Aconitase Activity Assay Kit | Sigma Aldrich | MAK051 | |
| Succinate Colorimetric Assay Kit | Sigma Aldrich | MAK184 | |
| Succinate Dehydrogenase Activity Colorimetric Assay Kit | Sigma Aldrich | MAK197 | |
| QIAprep® Spin Miniprep Kit | Qiagen | 27106 | |
| **Equipment** | | | |
| Model 120 Sonic Dismembrator | Fisherbrand | FB120110 | |
| CFX96 Touch Real-Time PCR Detection System | Bio-rad | Discontinued | |

| | | |
|---|---|---|
| C1000 Touch Thermocycler | Bio-rad | 1851148 |
| Nanoject III | Drummond Scientific Company | 3-000-207 |
| Cytospin 4 | Thermo Scientific | A78300003 |
| **Cell lines** | | |
| *Ixodes scapularis* ISE6 cells | Ulrike Munderloh, University of Minnesota | ISE6 |
| *Ixodes* scapularis IDE12 cells | Ulrike Munderloh, University of Minnesota | IDE12 |
| *Amblyomma americanum* AAE2 cells | Ulrike Munderloh, University of Minnesota | AAE2 |
| *Dermacentor andersoni* DAE100 cells | Ulrike Munderloh, University of Minnesota | DAE100 |
| HL-60 | ATCC | CCL-240 |
| **Organisms** | | |
| *Ixodes scapularis* Nymphs | Tick Lab, Oklahoma State University | N/A |
| *Ixodes scapularis* Nymphs | Ulrike Munderloh, University of Minnesota | N/A |
| C57BL6J Mice | University of Maryland, Baltimore | N/A |
| C57BL6J Mice | Jackson Laboratories | #000664 |
| *Borrelia burgdorferi* B31 clone MSK5 | Jon Skare, Texas A&M University | N/A |
| *Anaplasma phagocytophilum* HZ | Ulrike Munderloh, University of Minnesota | N/A |
| *Rickettsia buchneri* ISO7$^{T}$ | Ulrike Munderloh, University of Minnesota | N/A |

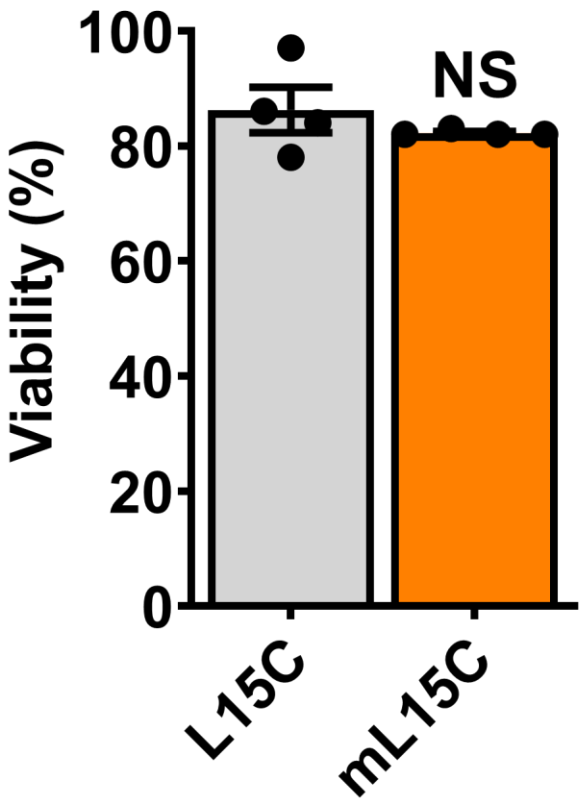

**Figure S1**

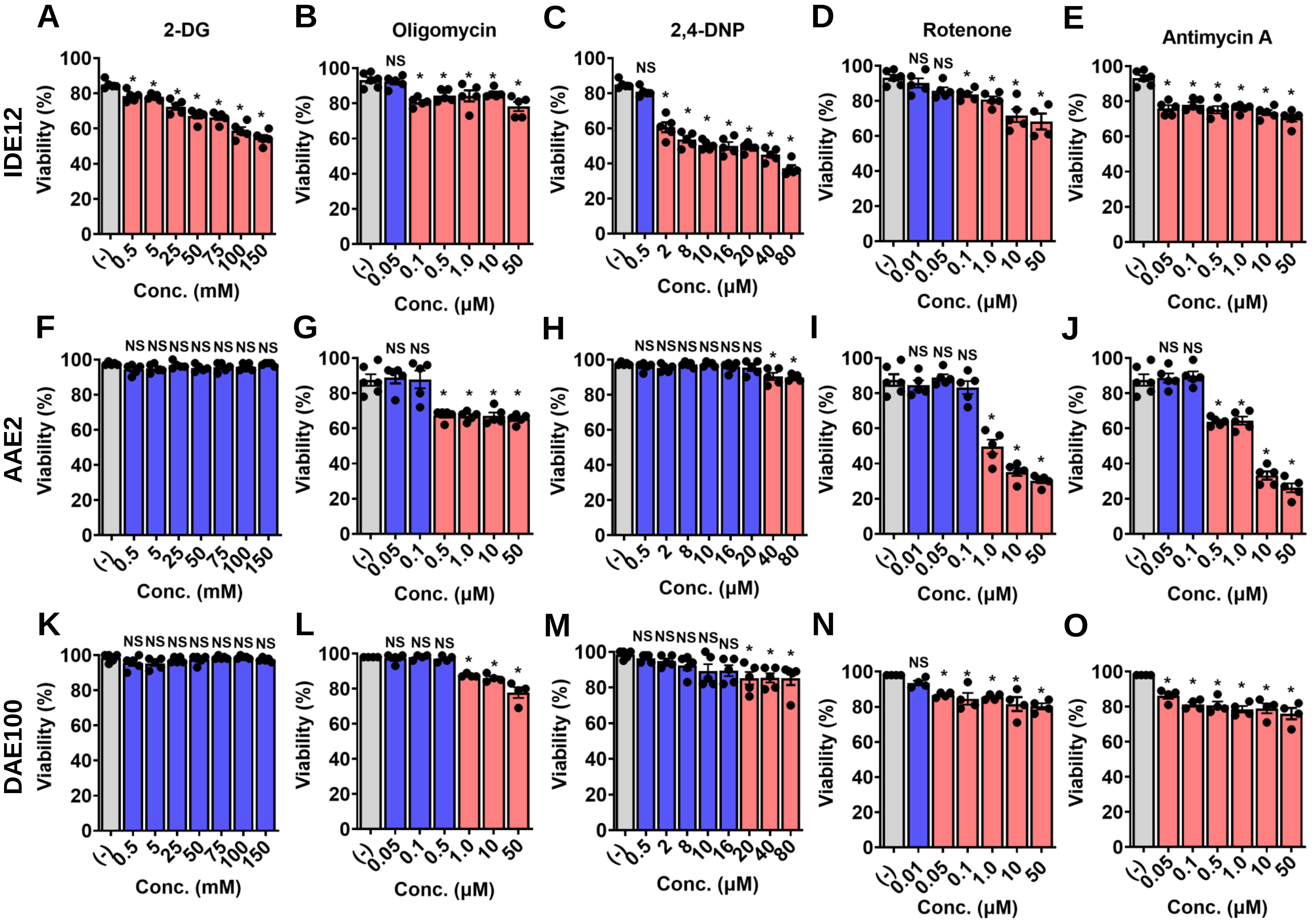
A
2-DG
B
Oligomycin
C
2,4-DNP
D
Rotenone
E
Antimycin A
IDE12
F
G
H
I
J
AAE2
K
L
M
N
O
DAE100
Viability (%)
Conc. (mM)
Conc. (μM)
NS
*

Figure S2

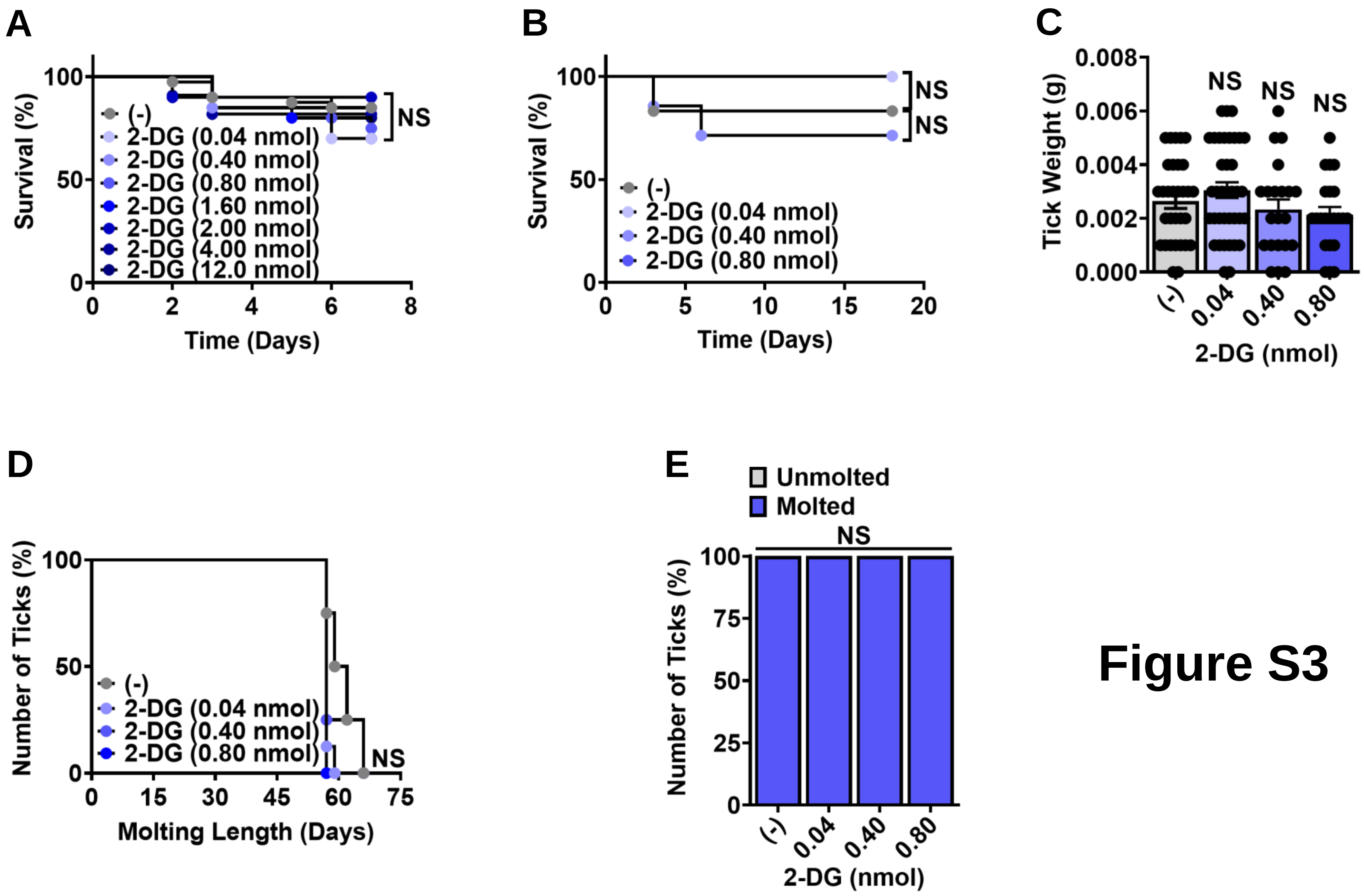
A
Survival (%)
Time (Days)
(-)
2-DG (0.04 nmol)
2-DG (0.40 nmol)
2-DG (0.80 nmol)
2-DG (1.60 nmol)
2-DG (2.00 nmol)
2-DG (4.00 nmol)
2-DG (12.0 nmol)
NS
B
Survival (%)
Time (Days)
(-)
2-DG (0.04 nmol)
2-DG (0.40 nmol)
2-DG (0.80 nmol)
NS
NS
C
Tick Weight (g)
NS
NS
NS
(-)
0.04
0.40
0.80
2-DG (nmol)
D
Number of Ticks (%)
Molting Length (Days)
(-)
2-DG (0.04 nmol)
2-DG (0.40 nmol)
2-DG (0.80 nmol)
NS
E
Unmolted
Molted
NS
Number of Ticks (%)
(-)
0.04
0.40
0.80
2-DG (nmol)

Figure S3

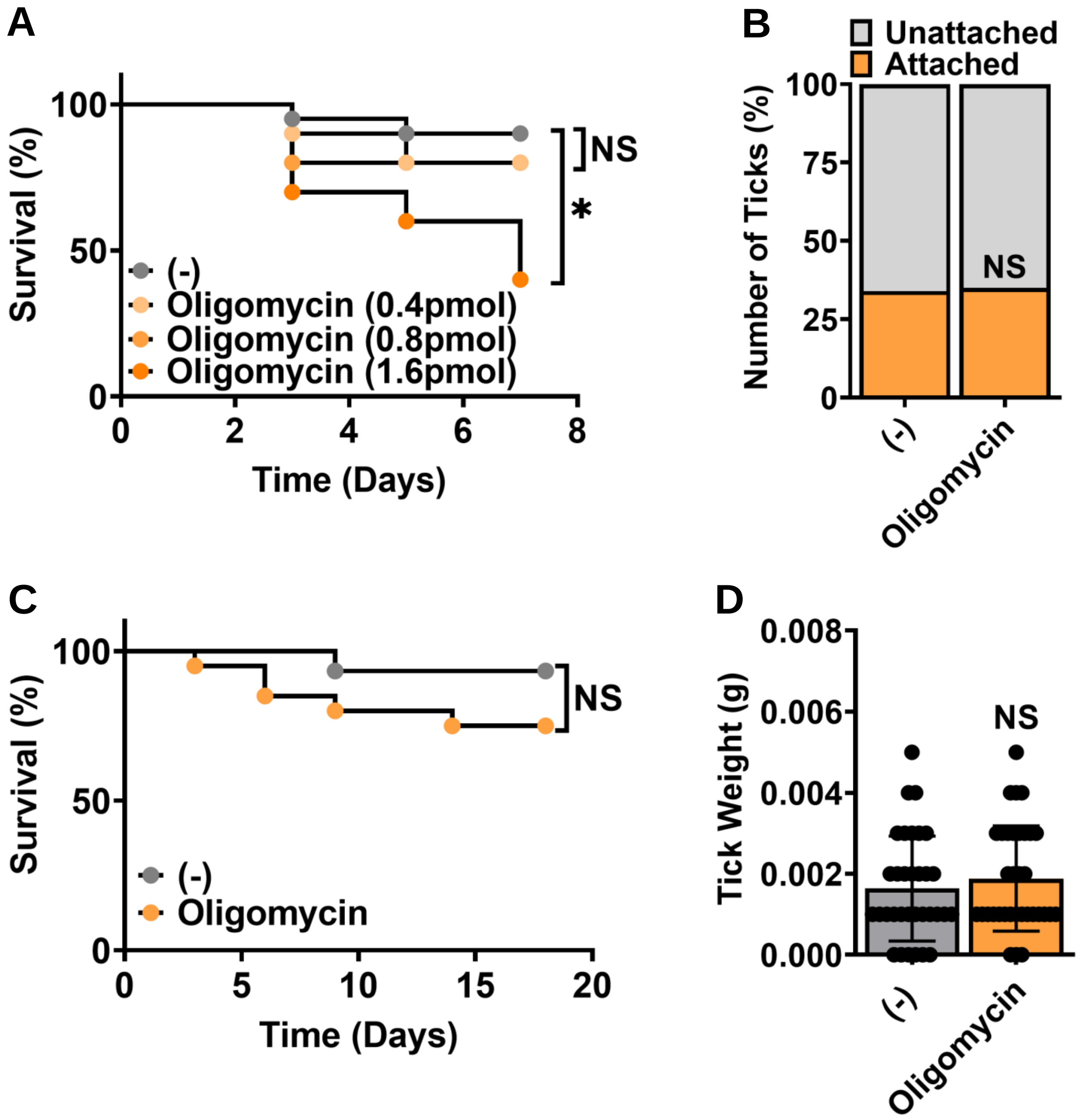
A
Survival (%)
100
50
0
0
2
4
6
8
Time (Days)
NS
*
(-)
Oligomycin (0.4pmol)
Oligomycin (0.8pmol)
Oligomycin (1.6pmol)
B
Unattached
Attached
Number of Ticks (%)
100
75
50
25
0
NS
(-)
Oligomycin
C
Survival (%)
100
50
0
0
5
10
15
20
Time (Days)
NS
(-)
Oligomycin
D
Tick Weight (g)
0.008
0.006
0.004
0.002
0.000
NS
(-)
Oligomycin

Figure S4

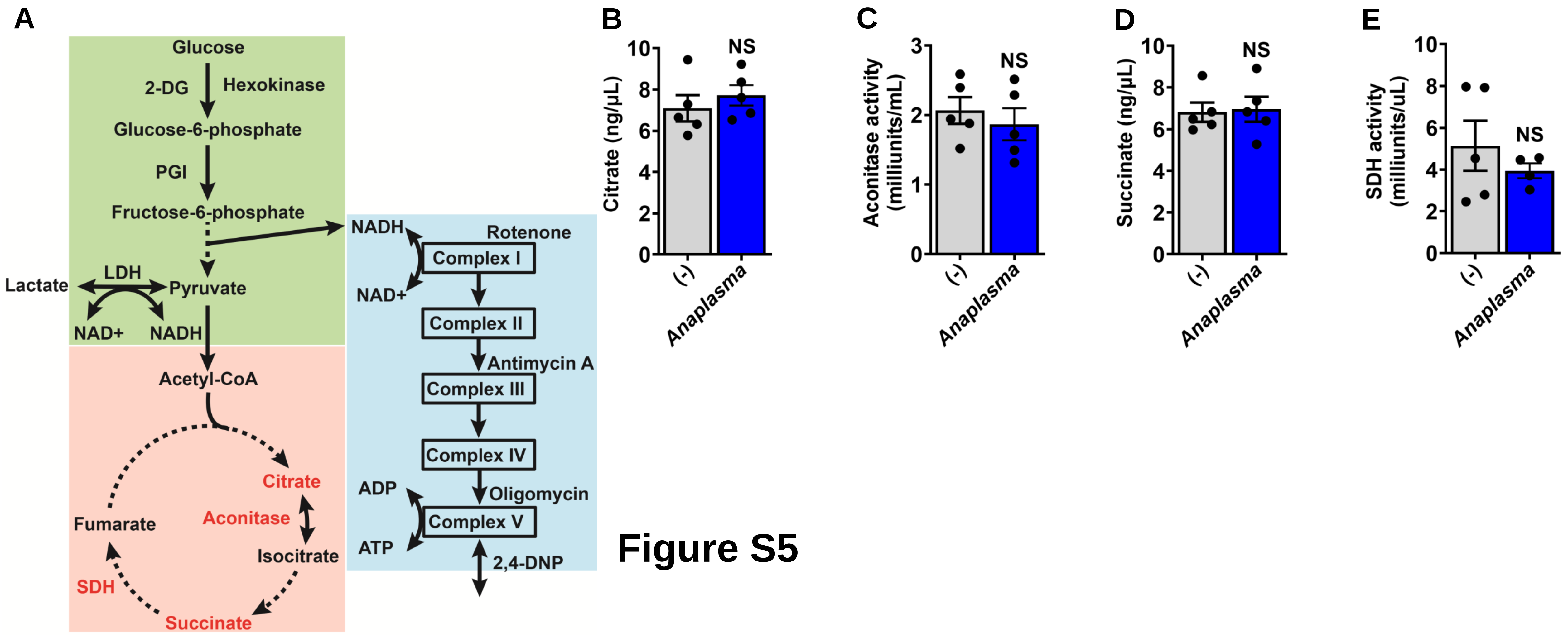
A
Glucose
2-DG
Hexokinase
Glucose-6-phosphate
PGI
Fructose-6-phosphate
LDH
Lactate
Pyruvate
NAD+
NADH
Acetyl-CoA
Citrate
Aconitase
Fumarate
Isocitrate
SDH
Succinate
NADH
Rotenone
Complex I
NAD+
Complex II
Antimycin A
Complex III
Complex IV
ADP
Oligomycin
Complex V
ATP
2,4-DNP
B
Citrate (ng/μL)
NS
(-)
Anaplasma
C
Aconitase activity (milliunits/mL)
NS
(-)
Anaplasma
D
Succinate (ng/μL)
NS
(-)
Anaplasma
E
SDH activity (milliunits/uL)
NS
(-)
Anaplasma

### Figure S5

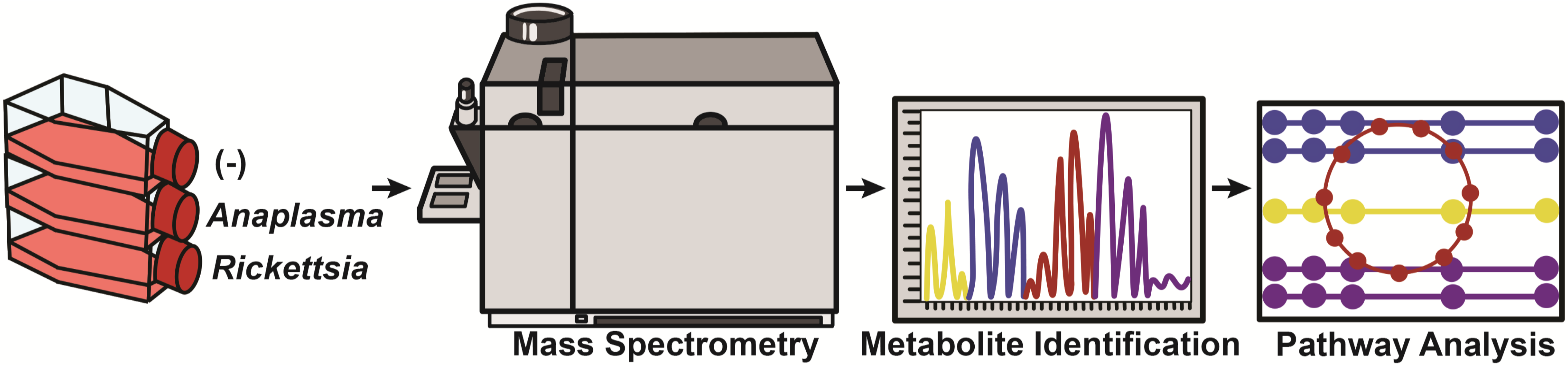
(-)
Anaplasma
Rickettsia
Mass Spectrometry
Metabolite Identification
Pathway Analysis

**Figure S6**

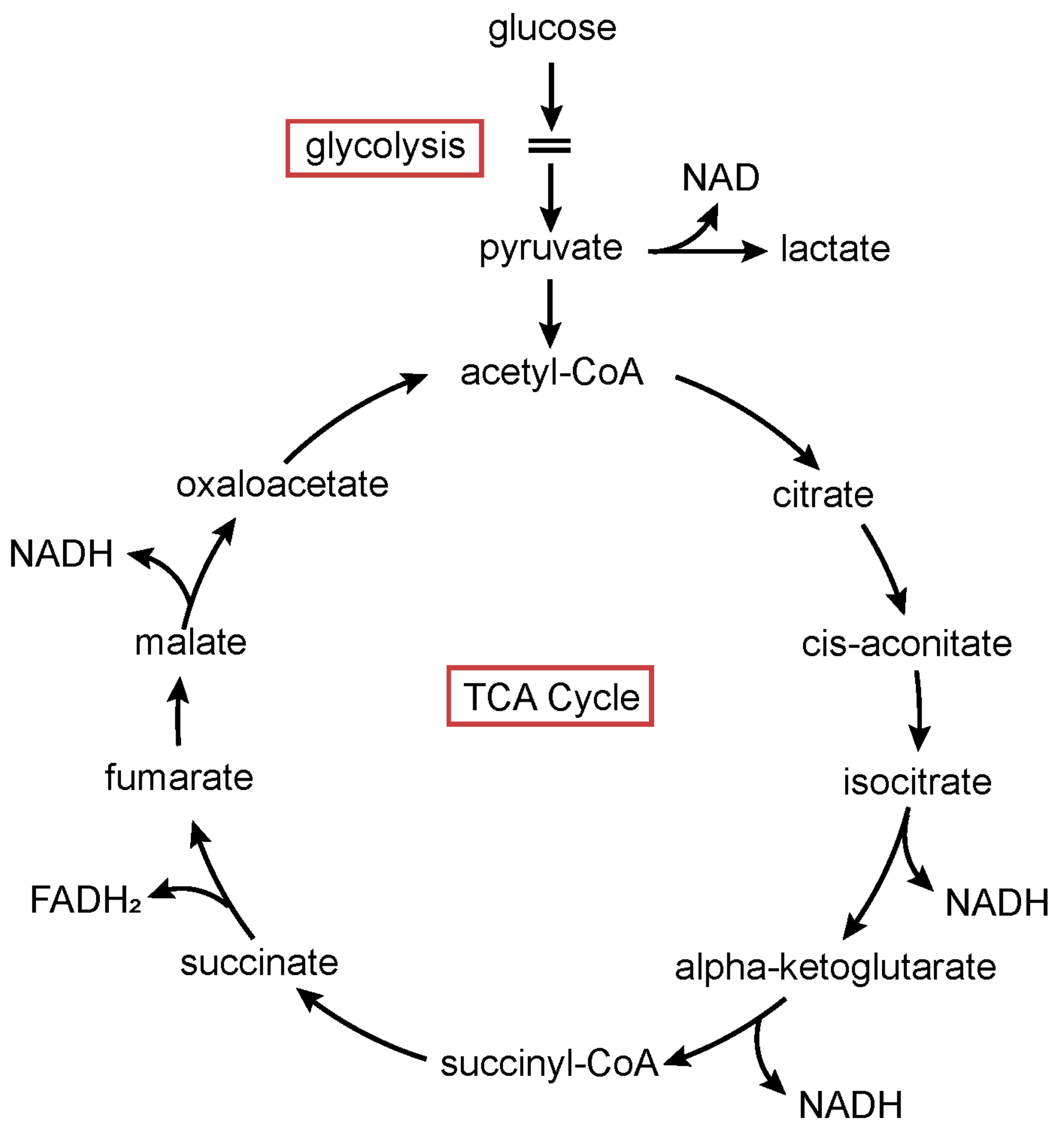
glucose
glycolysis
NAD
pyruvate
lactate
acetyl-CoA
citrate
cis-aconitate
isocitrate
NADH
alpha-ketoglutarate
NADH
succinyl-CoA
succinate
FADH₂
fumarate
malate
NADH
oxaloacetate
TCA Cycle

### Figure S7

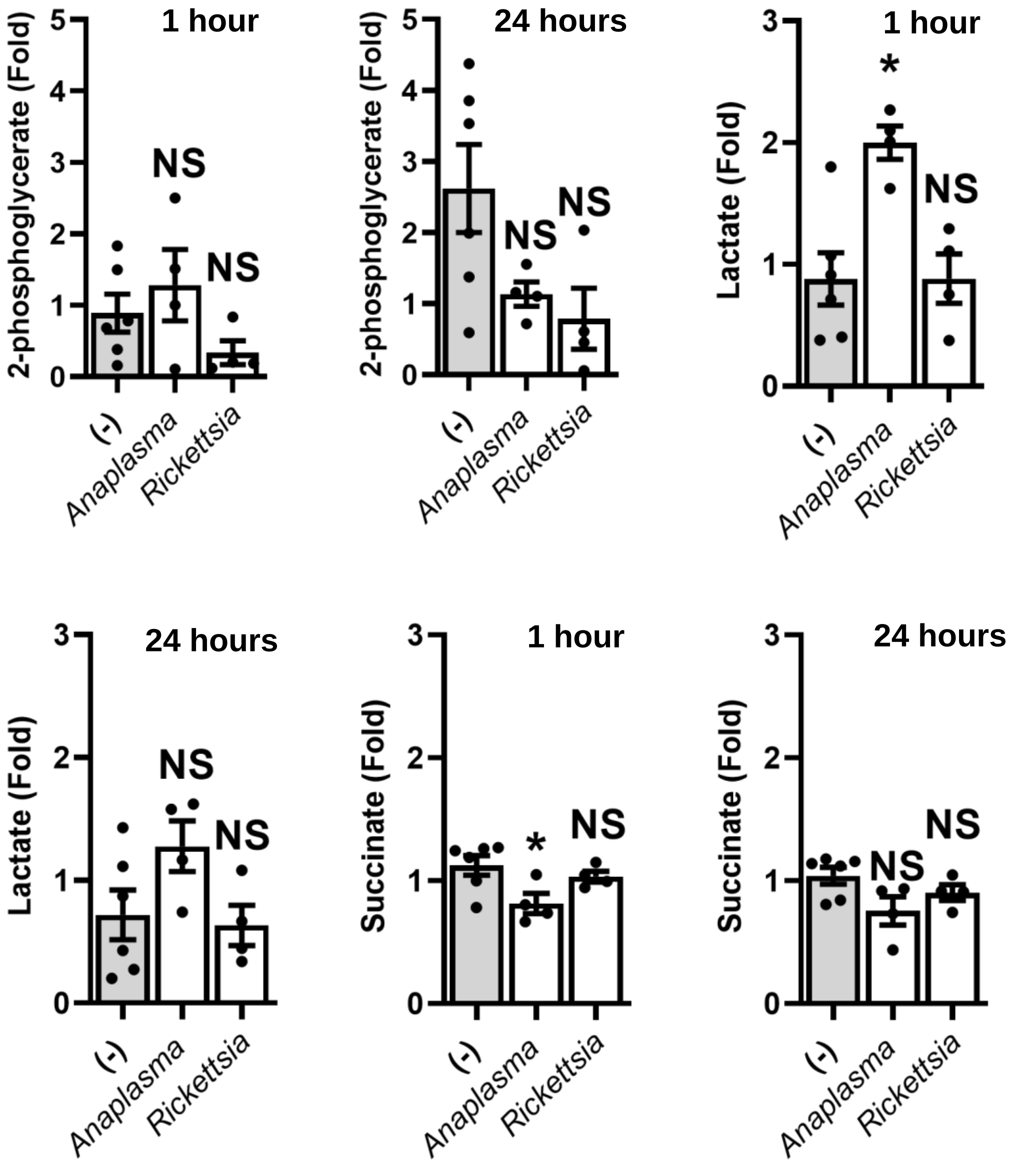
1 hour
24 hours
1 hour
24 hours
1 hour
24 hours
2-phosphoglycerate (Fold)
Lactate (Fold)
Succinate (Fold)
NS
*
(-)
Anaplasma
Rickettsia

### Figure S8

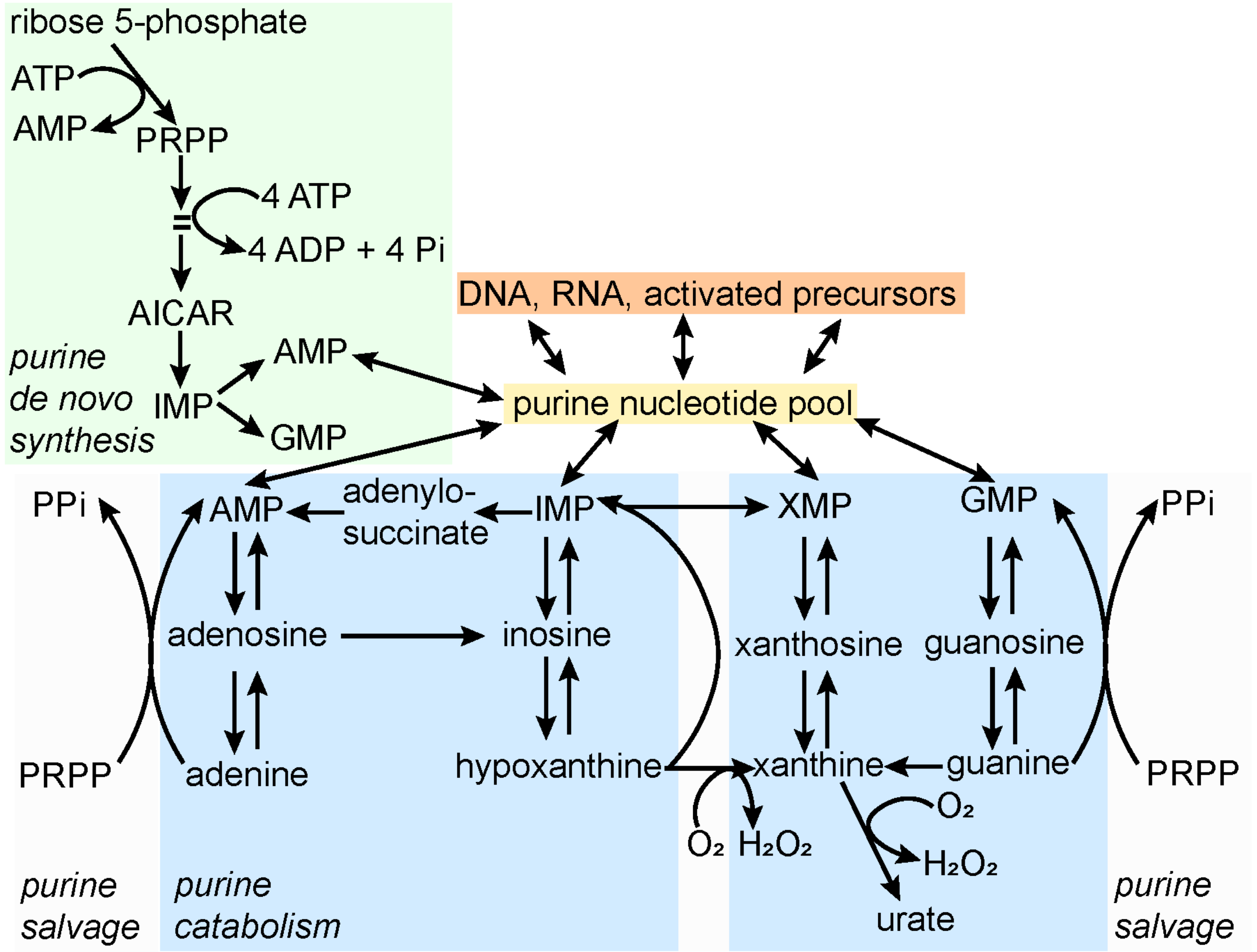
ribose 5-phosphate
ATP
AMP
PRPP
4 ATP
4 ADP + 4 Pi
AICAR
purine
de novo
synthesis
IMP
AMP
GMP
DNA, RNA, activated precursors
purine nucleotide pool
PPi
AMP
adenylo-
succinate
IMP
XMP
GMP
PPi
adenosine
inosine
xanthosine
guanosine
PRPP
adenine
hypoxanthine
xanthine
guanine
PRPP
O₂
H₂O₂
O₂
H₂O₂
urate
purine
salvage
purine
catabolism
purine
salvage

**Figure S9**

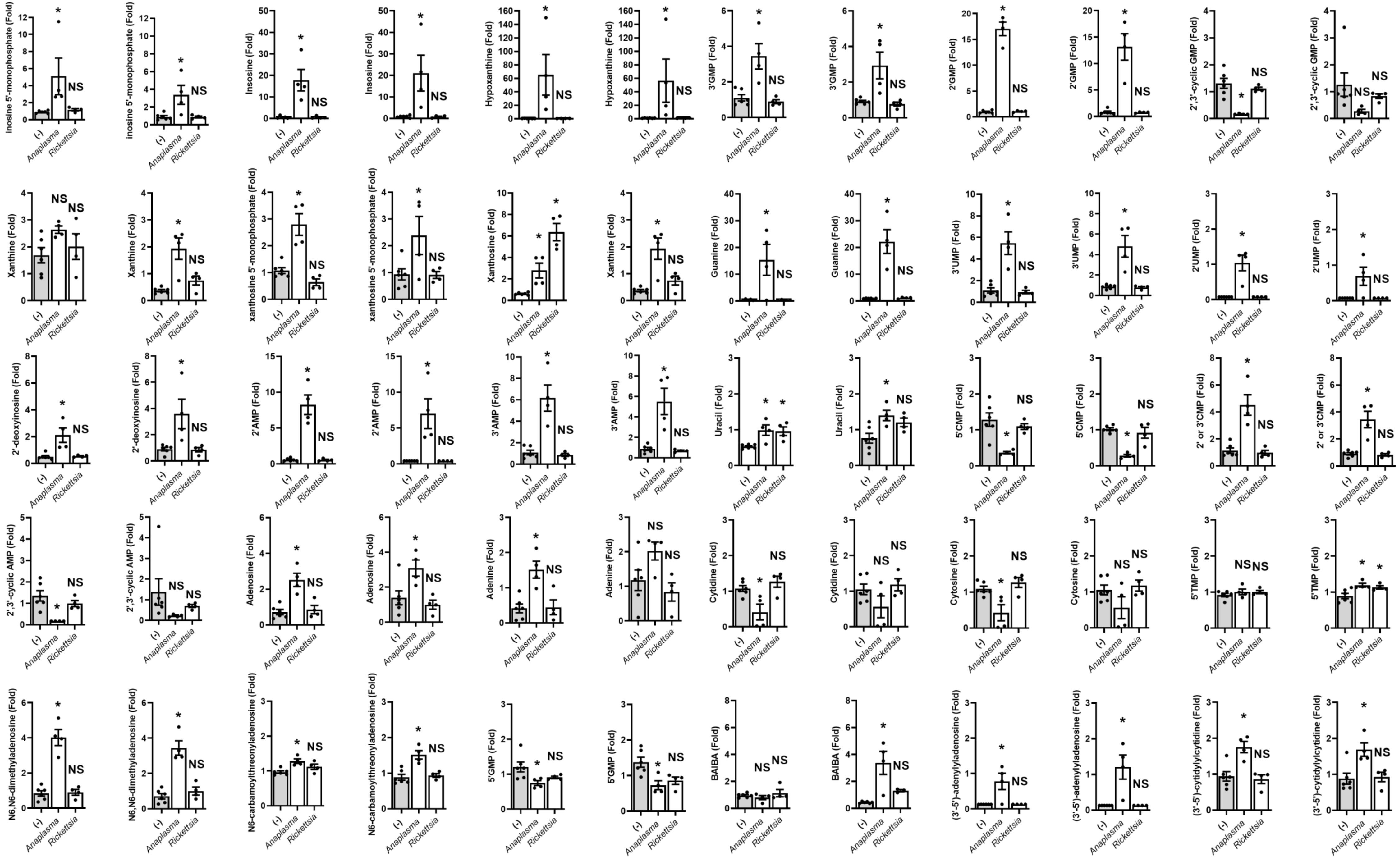

inosine 5'-monophosphate (Fold)
Insosine (Fold)
Inosine (Fold)
Hypoxanthine (Fold)
3'GMP (Fold)
2'GMP (Fold)
2',3'-cyclic GMP (Fold)
Xanthine (Fold)
xanthosine 5'-monophosphate (Fold)
Xanthosine (Fold)
Guanine (Fold)
3'UMP (Fold)
2'UMP (Fold)
2'-deoxyinosine (Fold)
2'AMP (Fold)
3'AMP (Fold)
Uracil (Fold)
5'CMP (Fold)
2' or 3'CMP (Fold)
2',3'-cyclic AMP (Fold)
Adenosine (Fold)
Adenine (Fold)
Cytidine (Fold)
Cytosine (Fold)
5'TMP (Fold)
N6,N6-dimethyladenosine (Fold)
N6-carbamoylthreonyladenosine (Fold)
5'GMP (Fold)
BAIBA (Fold)
(3'-5')-adenylyladenosine (Fold)
(3'-5')-cytidylylcytidine (Fold)
(-)
Anaplasma
Rickettsia
*
NS

### Figure S10

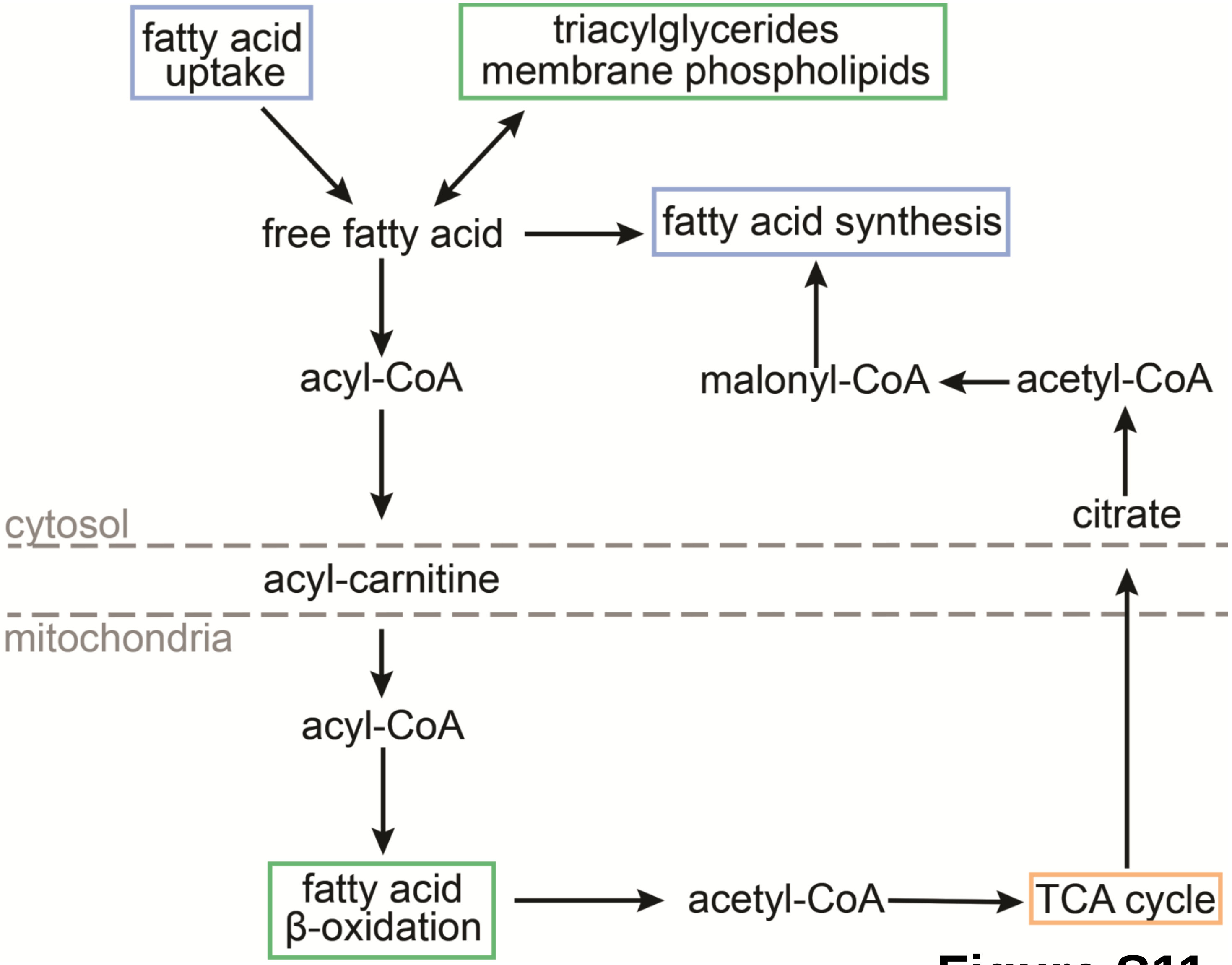
fatty acid uptake
triacylglycerides
membrane phospholipids
free fatty acid
fatty acid synthesis
acyl-CoA
malonyl-CoA
acetyl-CoA
cytosol
citrate
acyl-carnitine
mitochondria
acyl-CoA
fatty acid
β-oxidation
acetyl-CoA
TCA cycle

**Figure S11**

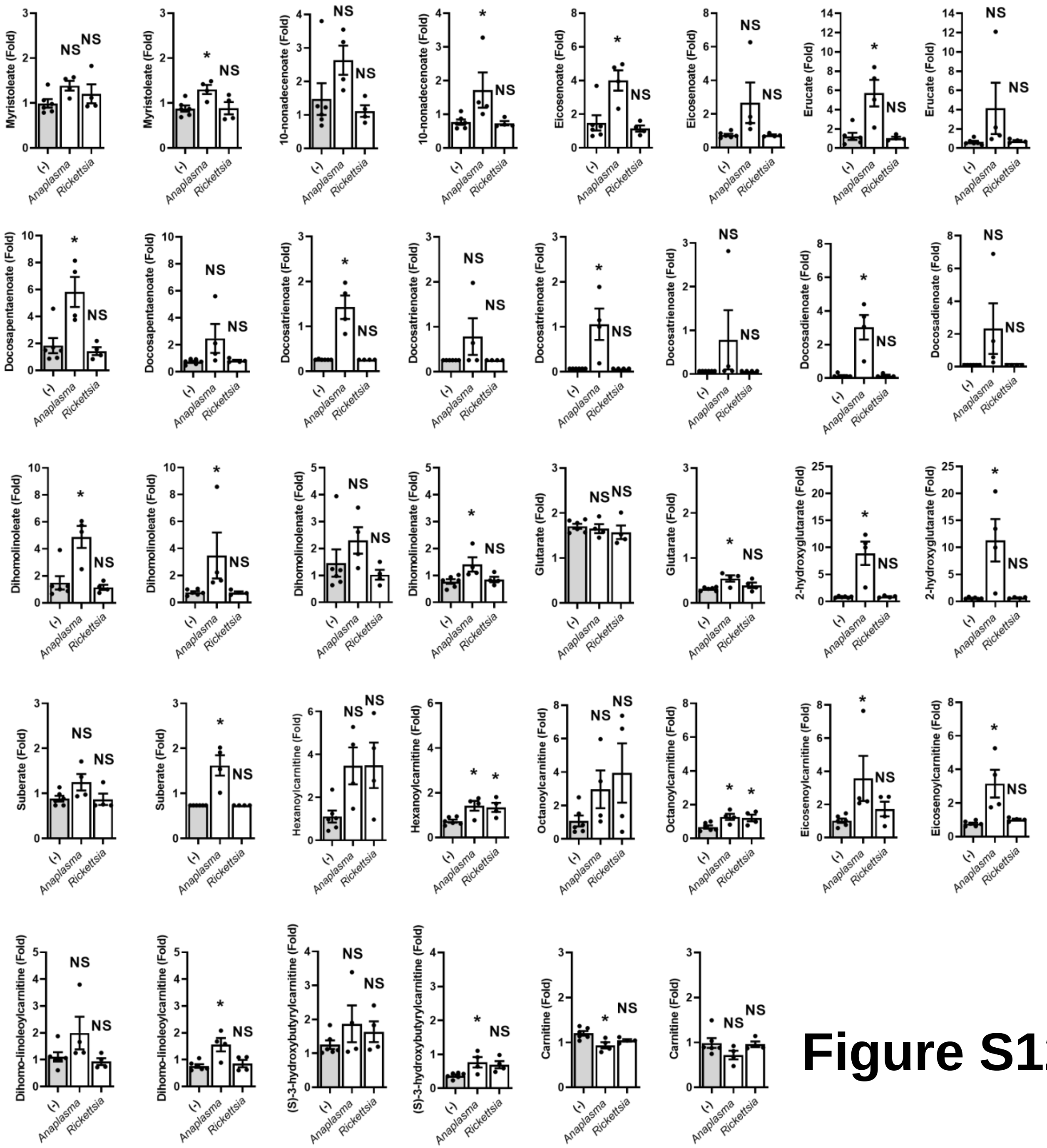

Myristoleate (Fold)
Myristoleate (Fold)
10-nonadecenoate (Fold)
10-nonadecenoate (Fold)
Eicosenoate (Fold)
Eicosenoate (Fold)
Erucate (Fold)
Erucate (Fold)
Docosapentaenoate (Fold)
Docosapentaenoate (Fold)
Docosatrienoate (Fold)
Docosatrienoate (Fold)
Docosatrienoate (Fold)
Docosatrienoate (Fold)
Docosadienoate (Fold)
Docosadienoate (Fold)
Dihomolinoleate (Fold)
Dihomolinoleate (Fold)
Dihomolinolenate (Fold)
Dihomolinolenate (Fold)
Glutarate (Fold)
Glutarate (Fold)
2-hydroxyglutarate (Fold)
2-hydroxyglutarate (Fold)
Suberate (Fold)
Suberate (Fold)
Hexanoylcarnitine (Fold)
Hexanoylcarnitine (Fold)
Octanoylcarnitine (Fold)
Octanoylcarnitine (Fold)
Eicosenoylcarnitine (Fold)
Eicosenoylcarnitine (Fold)
Dihomo-linoleoylcarnitine (Fold)
Dihomo-linoleoylcarnitine (Fold)
(S)-3-hydroxybutyrylcarnitine (Fold)
(S)-3-hydroxybutyrylcarnitine (Fold)
Carnitine (Fold)
Carnitine (Fold)
(-)
Anaplasma
Rickettsia
NS
*

### Figure S12

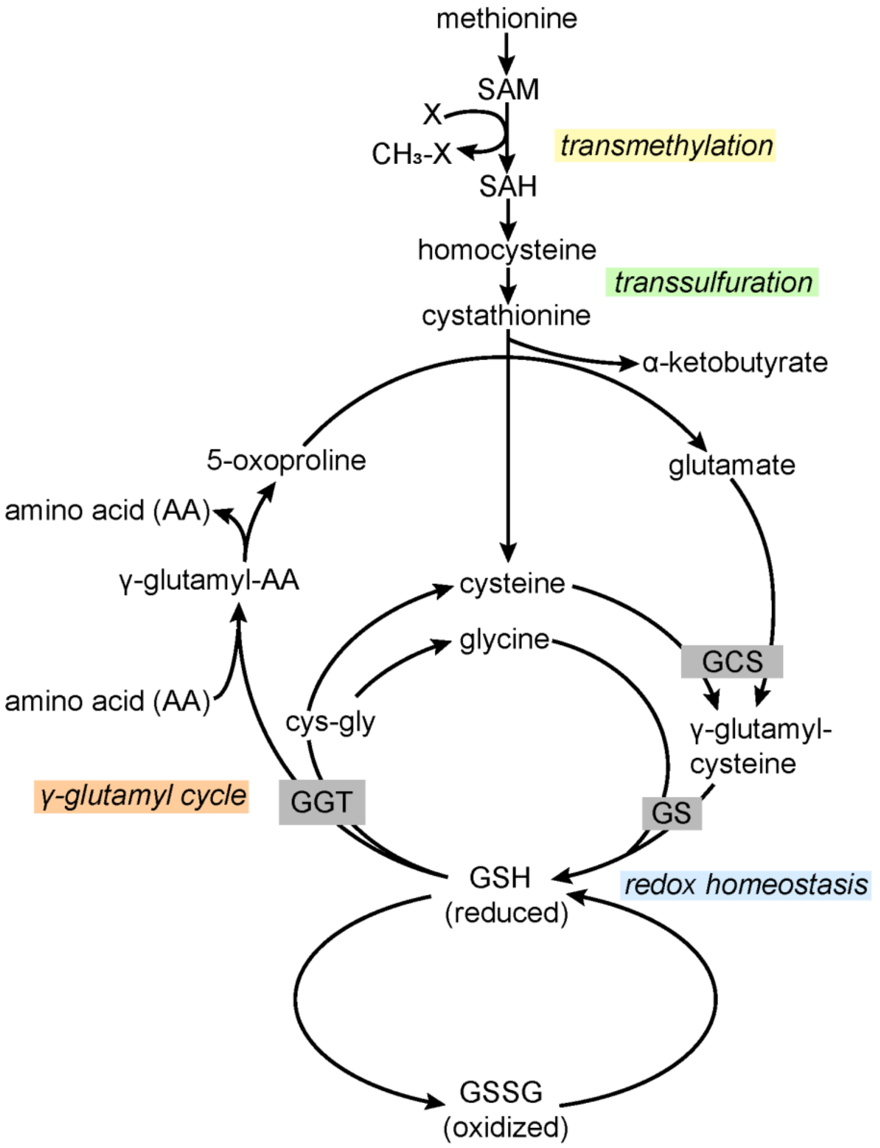
methionine
SAM
X
CH₃-X
SAH
transmethylation
homocysteine
transsulfuration
cystathionine
α-ketobutyrate
5-oxoproline
glutamate
amino acid (AA)
γ-glutamyl-AA
cysteine
glycine
GCS
amino acid (AA)
cys-gly
γ-glutamyl-
cysteine
γ-glutamyl cycle
GGT
GS
GSH
(reduced)
redox homeostasis
GSSG
(oxidized)

**Figure S13**

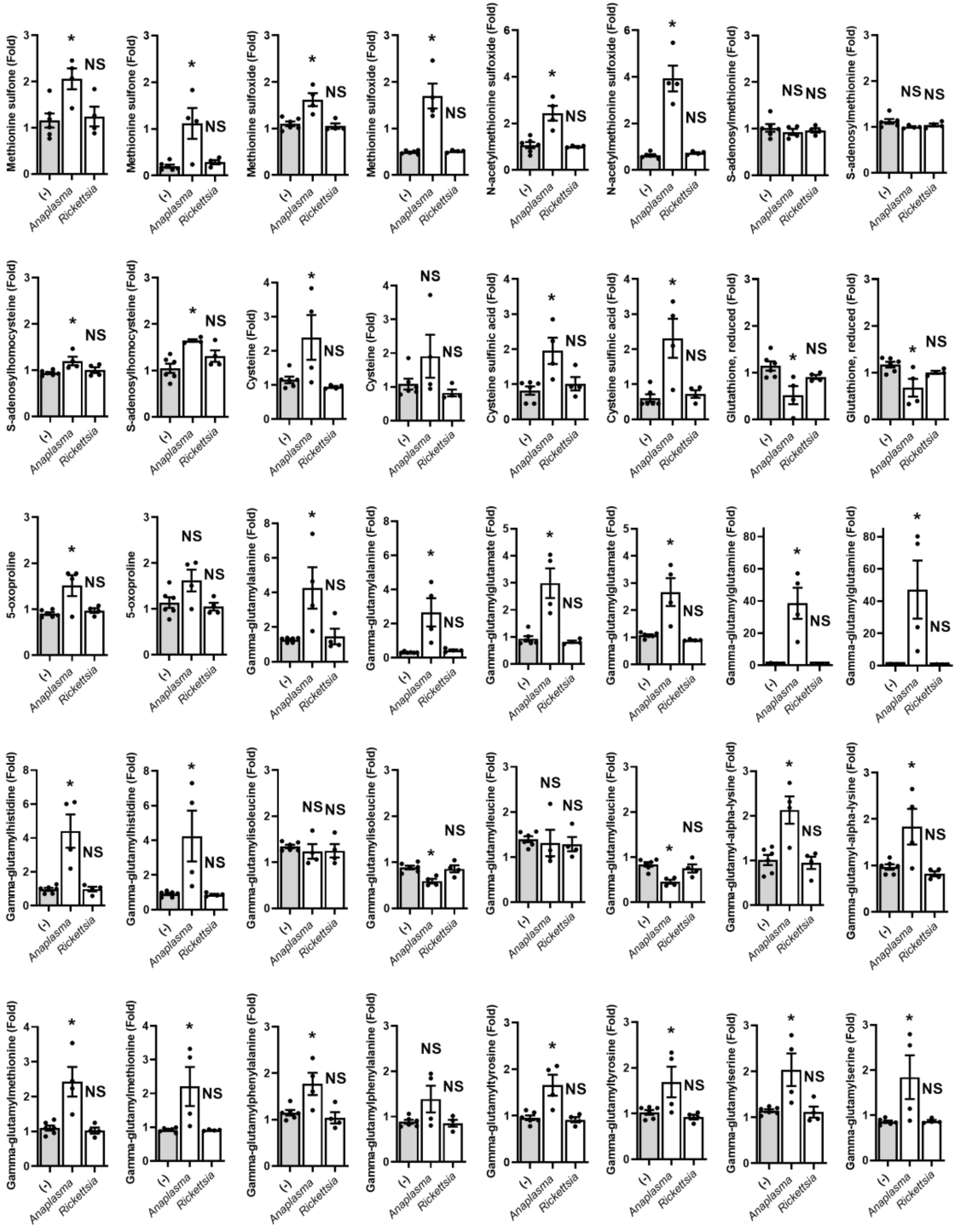
Methionine sulfone (Fold)
Methionine sulfone (Fold)
Methionine sulfoxide (Fold)
Methionine sulfoxide (Fold)
N-acetylmethionine sulfoxide (Fold)
N-acetylmethionine sulfoxide (Fold)
S-adenosylmethionine (Fold)
S-adenosylmethionine (Fold)
S-adenosylhomocysteine (Fold)
S-adenosylhomocysteine (Fold)
Cysteine (Fold)
Cysteine (Fold)
Cysteine sulfinic acid (Fold)
Cysteine sulfinic acid (Fold)
Glutathione, reduced (Fold)
Glutathione, reduced (Fold)
5-oxoproline
5-oxoproline
Gamma-glutamylalanine (Fold)
Gamma-glutamylalanine (Fold)
Gamma-glutamylglutamate (Fold)
Gamma-glutamylglutamate (Fold)
Gamma-glutamylglutamine (Fold)
Gamma-glutamylglutamine (Fold)
Gamma-glutamylhistidine (Fold)
Gamma-glutamylhistidine (Fold)
Gamma-glutamylisoleucine (Fold)
Gamma-glutamylisoleucine (Fold)
Gamma-glutamylleucine (Fold)
Gamma-glutamylleucine (Fold)
Gamma-glutamyl-alpha-lysine (Fold)
Gamma-glutamyl-alpha-lysine (Fold)
Gamma-glutamylmethionine (Fold)
Gamma-glutamylmethionine (Fold)
Gamma-glutamylphenylalanine (Fold)
Gamma-glutamylphenylalanine (Fold)
Gamma-glutamyltyrosine (Fold)
Gamma-glutamyltyrosine (Fold)
Gamma-glutamylserine (Fold)
Gamma-glutamylserine (Fold)
(-)
*Anaplasma*
*Rickettsia*
NS
*

**Figure S14**

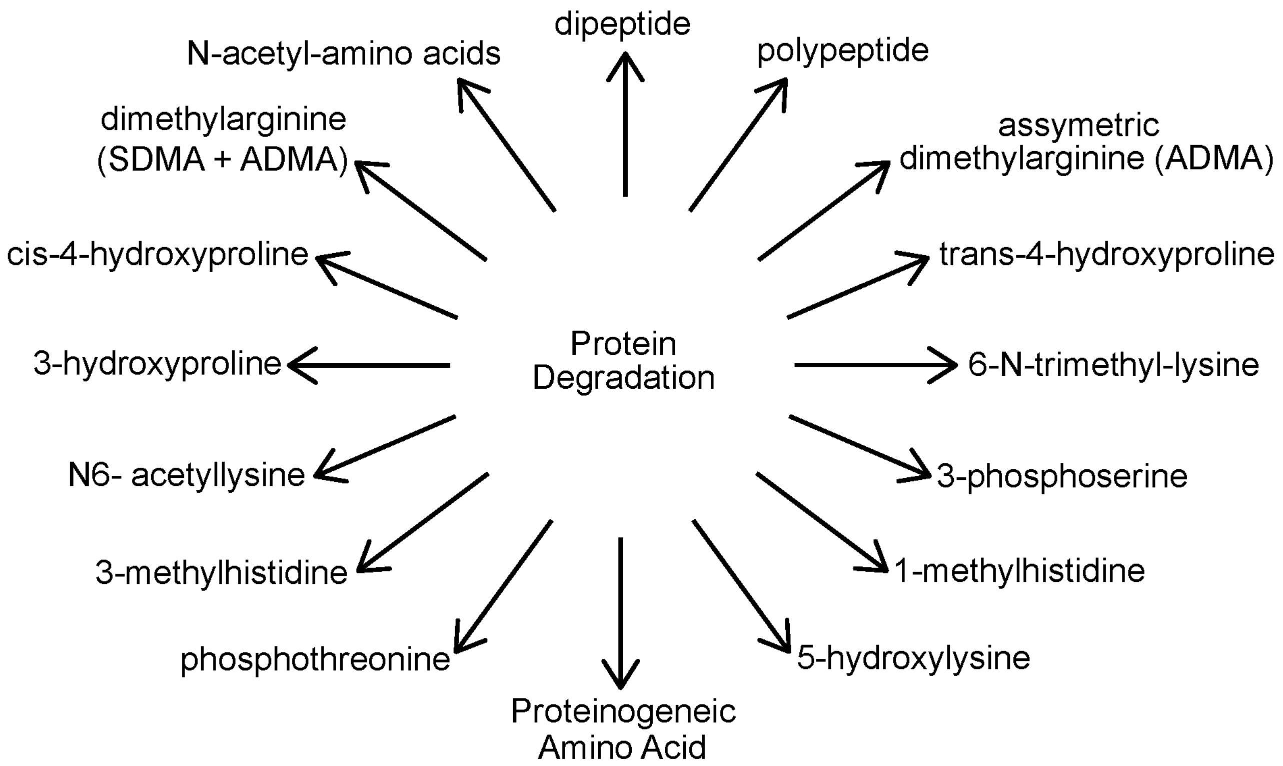
dipeptide
N-acetyl-amino acids
polypeptide
dimethylarginine
(SDMA + ADMA)
assymetric
dimethylarginine (ADMA)
cis-4-hydroxyproline
trans-4-hydroxyproline
Protein
Degradation
3-hydroxyproline
6-N-trimethyl-lysine
N6- acetyllysine
3-phosphoserine
3-methylhistidine
1-methylhistidine
phosphothreonine
5-hydroxylysine
Proteinogeneic
Amino Acid

**Figure S15**

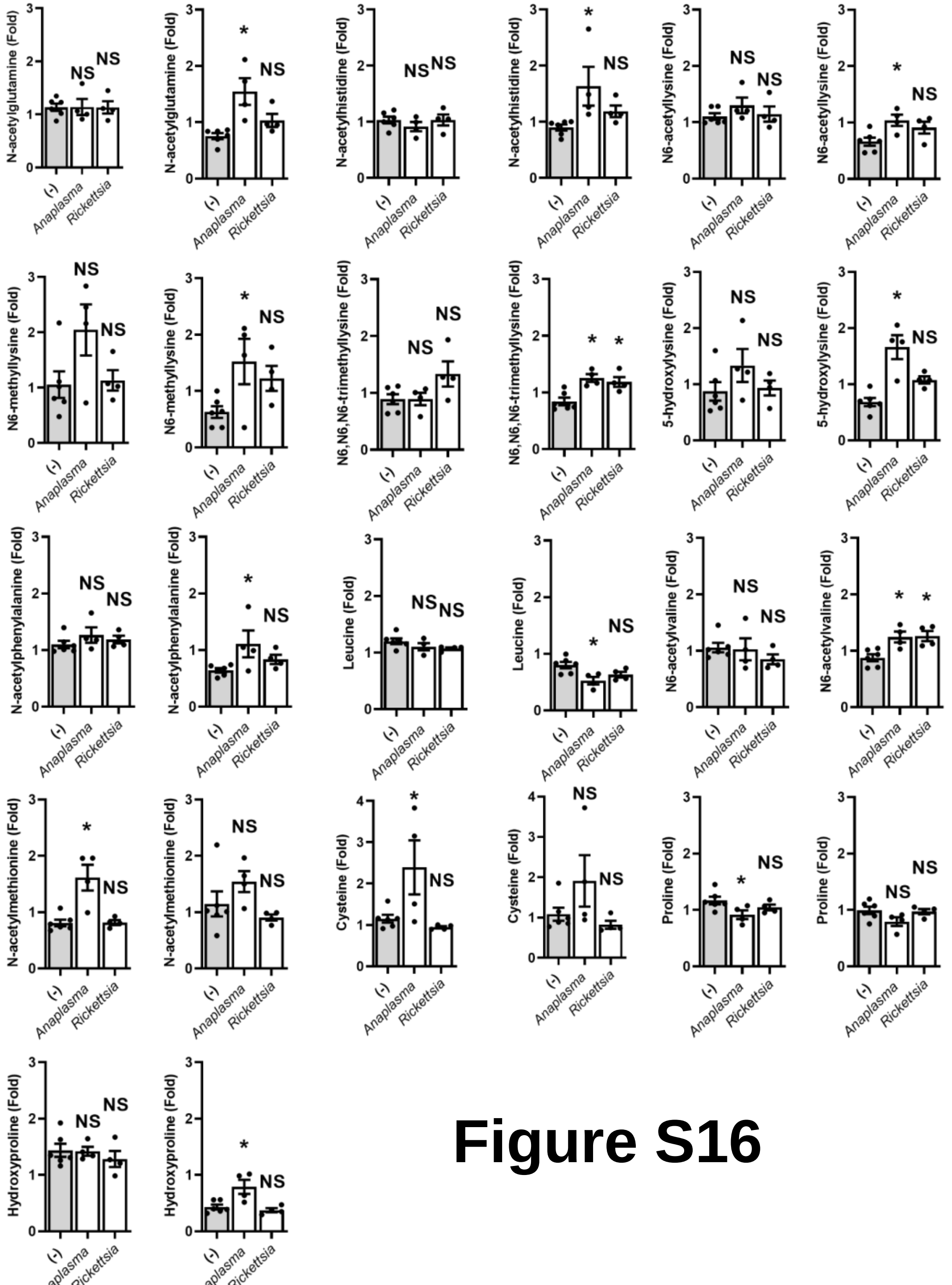
N-acetylglutamine (Fold)
N-acetylhistidine (Fold)
N6-acetyllysine (Fold)
N6-methyllysine (Fold)
N6,N6,N6-trimethyllysine (Fold)
5-hydroxylysine (Fold)
N-acetylphenylalanine (Fold)
Leucine (Fold)
N6-acetylvaline (Fold)
N-acetylmethionine (Fold)
Cysteine (Fold)
Proline (Fold)
Hydroxyproline (Fold)
(-)
Anaplasma
Rickettsia
NS
*

### Figure S16

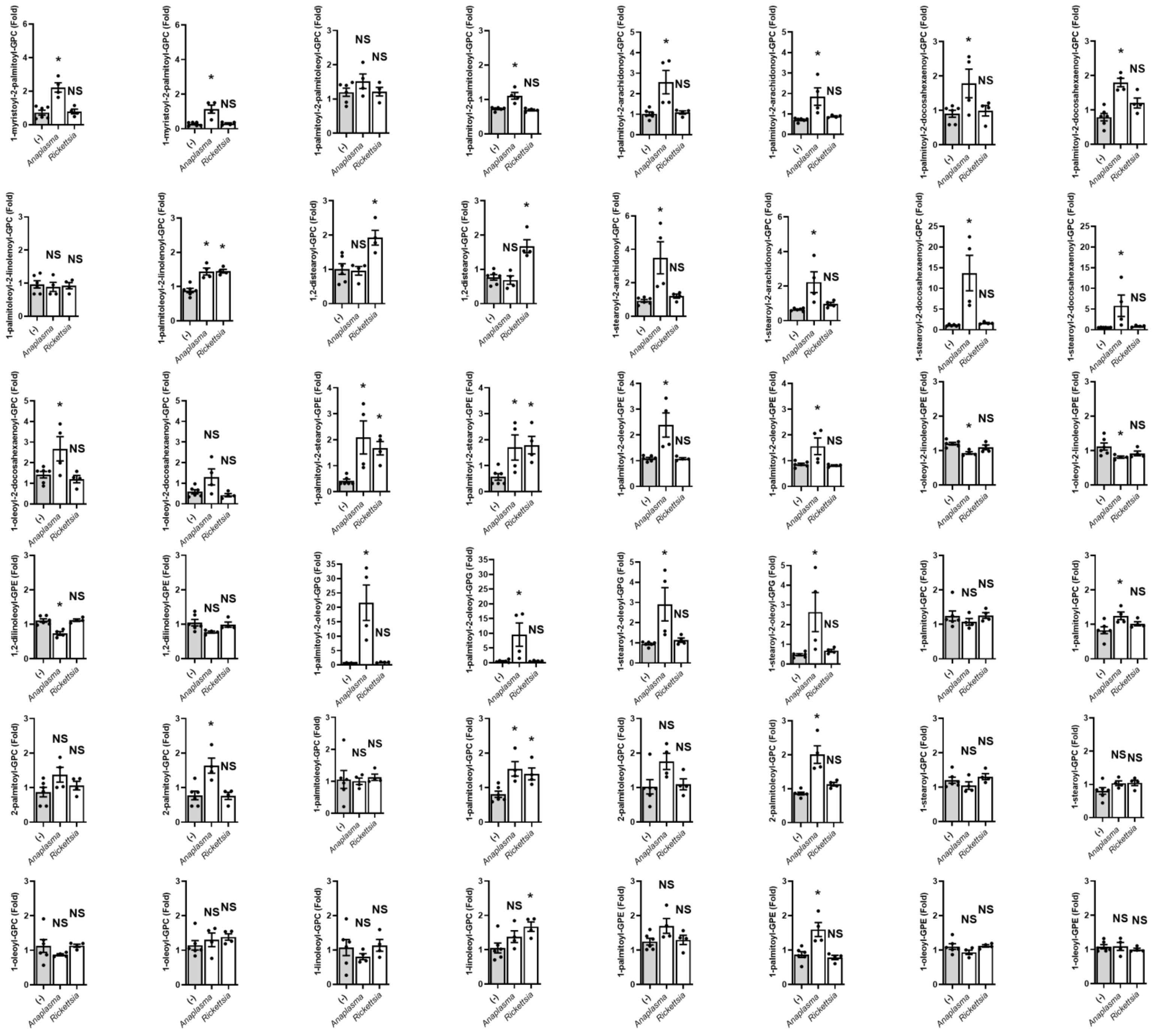
1-myristoyl-2-palmitoyl-GPC (Fold)
1-myristoyl-2-palmitoyl-GPC (Fold)
1-palmitoyl-2-palmitoleoyl-GPC (Fold)
1-palmitoyl-2-palmitoleoyl-GPC (Fold)
1-palmitoyl-2-arachidonoyl-GPC (Fold)
1-palmitoyl-2-arachidonoyl-GPC (Fold)
1-palmitoyl-2-docosahexaenoyl-GPC (Fold)
1-palmitoyl-2-docosahexaenoyl-GPC (Fold)
1-palmitoleoyl-2-linolenoyl-GPC (Fold)
1-palmitoleoyl-2-linolenoyl-GPC (Fold)
1,2-distearoyl-GPC (Fold)
1,2-distearoyl-GPC (Fold)
1-stearoyl-2-arachidonoyl-GPC (Fold)
1-stearoyl-2-arachidonoyl-GPC (Fold)
1-stearoyl-2-docosahexaenoyl-GPC (Fold)
1-stearoyl-2-docosahexaenoyl-GPC (Fold)
1-oleoyl-2-docosahexaenoyl-GPC (Fold)
1-oleoyl-2-docosahexaenoyl-GPC (Fold)
1-palmitoyl-2-stearoyl-GPE (Fold)
1-palmitoyl-2-stearoyl-GPE (Fold)
1-palmitoyl-2-oleoyl-GPE (Fold)
1-palmitoyl-2-oleoyl-GPE (Fold)
1-oleoyl-2-linoleoyl-GPE (Fold)
1-oleoyl-2-linoleoyl-GPE (Fold)
1,2-dilinoleoyl-GPE (Fold)
1,2-dilinoleoyl-GPE (Fold)
1-palmitoyl-2-oleoyl-GPG (Fold)
1-palmitoyl-2-oleoyl-GPG (Fold)
1-stearoyl-2-oleoyl-GPG (Fold)
1-stearoyl-2-oleoyl-GPG (Fold)
1-palmitoyl-GPC (Fold)
1-palmitoyl-GPC (Fold)
2-palmitoyl-GPC (Fold)
2-palmitoyl-GPC (Fold)
1-palmitoleoyl-GPC (Fold)
1-palmitoleoyl-GPC (Fold)
2-palmitoleoyl-GPC (Fold)
2-palmitoleoyl-GPC (Fold)
1-stearoyl-GPC (Fold)
1-stearoyl-GPC (Fold)
1-oleoyl-GPC (Fold)
1-oleoyl-GPC (Fold)
1-linoleoyl-GPC (Fold)
1-linoleoyl-GPC (Fold)
1-palmitoyl-GPE (Fold)
1-palmitoyl-GPE (Fold)
1-oleoyl-GPE (Fold)
1-oleoyl-GPE (Fold)
(-)
Anaplasma
Rickettsia
NS
*

**Figure S17**

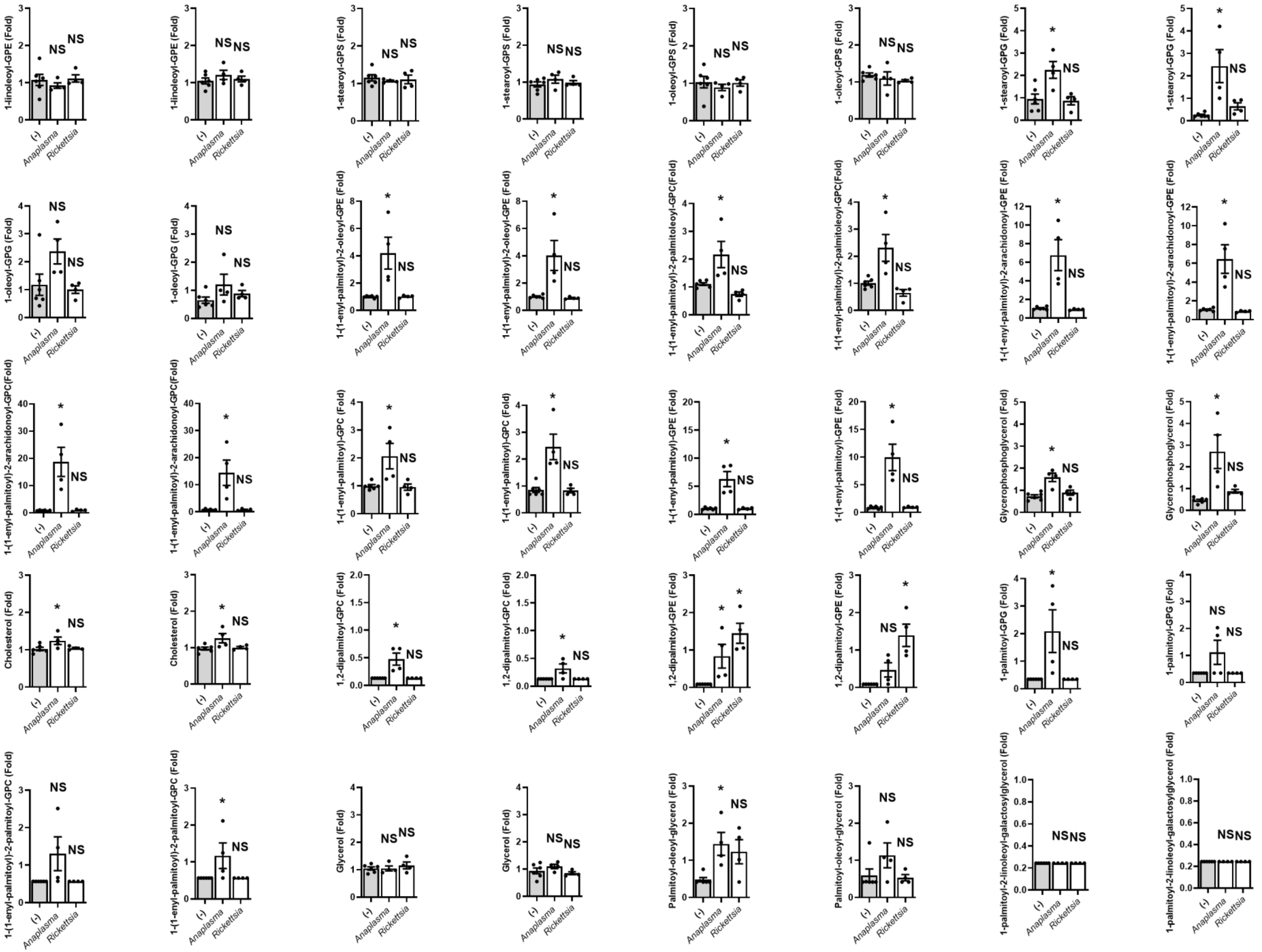

**Figure S17**
(*continuation*)

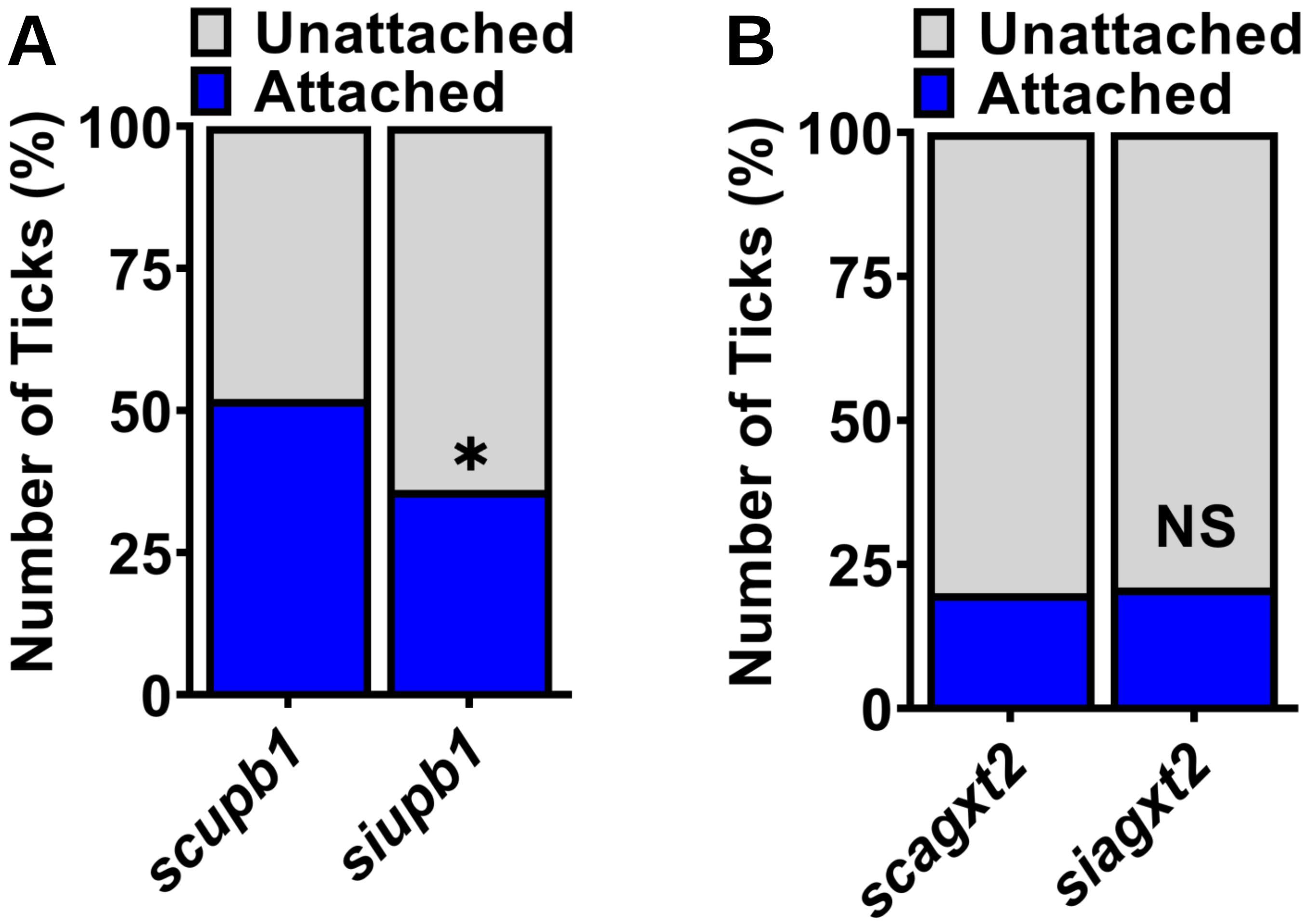
A
Unattached
Attached
Number of Ticks (%)
100
75
50
25
0
*
scupb1
siupb1
B
Unattached
Attached
Number of Ticks (%)
100
75
50
25
0
NS
scagxt2
siagxt2

Figure S18

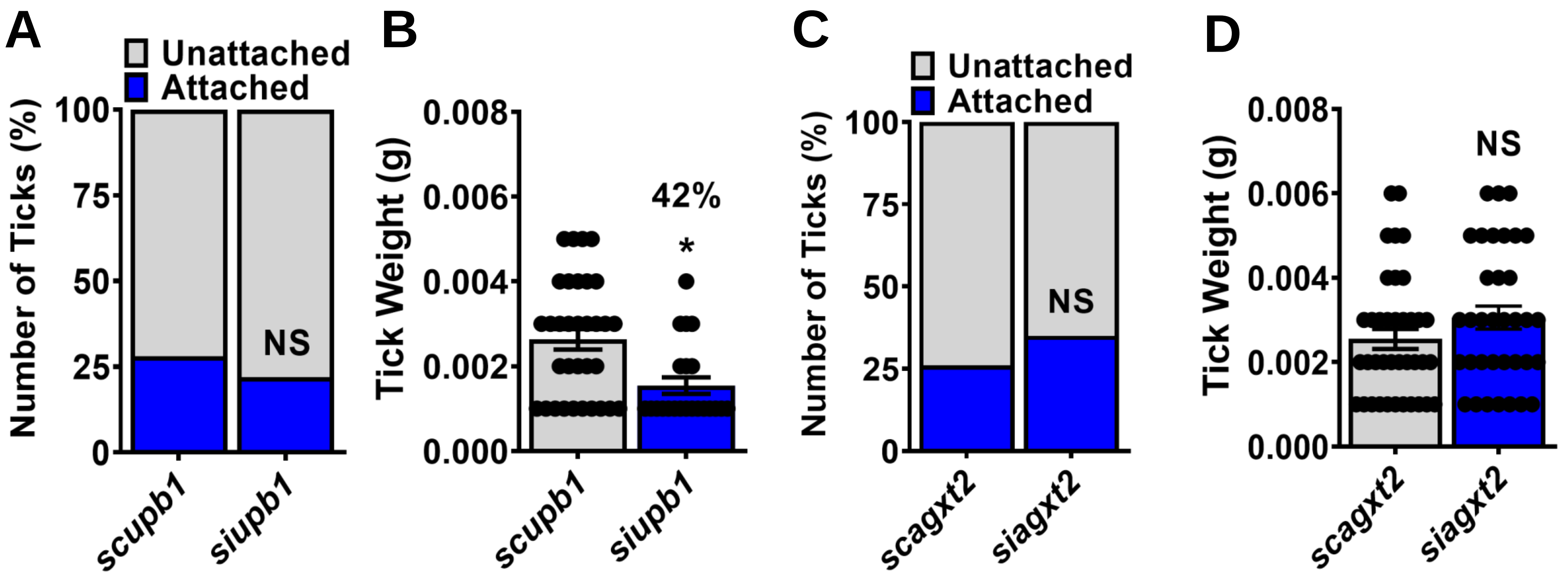
A
Unattached
Attached
Number of Ticks (%)
100
75
50
25
0
NS
scupb1
siupb1
B
Tick Weight (g)
0.008
0.006
0.004
0.002
0.000
42%
*
scupb1
siupb1
C
Unattached
Attached
Number of Ticks (%)
100
75
50
25
0
NS
scagxt2
siagxt2
D
Tick Weight (g)
0.008
0.006
0.004
0.002
0.000
NS
scagxt2
siagxt2

**Figure S19**

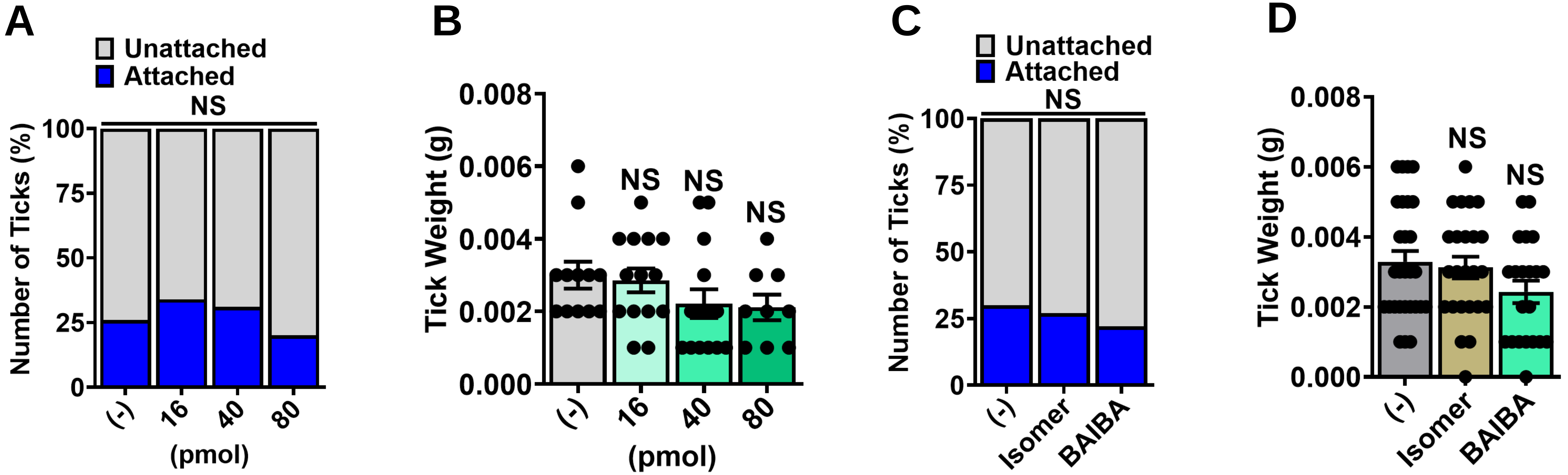
A
Unattached
Attached
NS
Number of Ticks (%)
100
75
50
25
0
(-)
16
40
80
(pmol)
B
Tick Weight (g)
0.008
0.006
0.004
0.002
0.000
NS
NS
NS
(-)
16
40
80
(pmol)
C
Unattached
Attached
NS
Number of Ticks (%)
100
75
50
25
0
(-)
Isomer
BAIBA
D
Tick Weight (g)
0.008
0.006
0.004
0.002
0.000
NS
NS
(-)
Isomer
BAIBA

Figure S20
